# GPU-Accelerated All-atom Particle-Mesh Ewald Continuous Constant pH Molecular Dynamics in Amber

**DOI:** 10.1101/2022.06.04.494833

**Authors:** Julie A. Harris, Ruibin Liu, Vinicius Martins de Oliveira, Erik Vaquez Montelongo, Jack A. Henderson, Jana Shen

## Abstract

Constant pH molecular dynamics (MD) simulations sample protonation states on the fly according to the conformational environment and user specified pH condition; however, the current accuracy is limited due to the use of implicit-solvent models or a hybrid solvent scheme. Here we report the first GPU-accelerated implementation, parameterization, and validation of the all-atom continuous constant pH MD (CpHMD) method with particle-mesh Ewald (PME) electrostatics in the Amber22 *pmemd. cuda* engine. The titration parameters for Asp, Glu, His, Cys, and Lys were derived for the CHARMM c22 and Amber ff14sb and ff19sb force fields. We then evaluated the PME-CpHMD method using the asynchronous pH replica-exchange titration simulations with the c22 force field for six benchmark proteins, including BBL, hen egg white lysozyme (HEWL), staphylococcal nuclease (SNase), thioredoxin, ribonuclease A (RNaseA), and human muscle creatine kinase (HMCK). The root-mean-square deviation from the experimental p*K*_a_’s of Asp, Glu, His, and Cys is 0.76 pH units, and the Pearson’s correlation coefficient for the p*K*_a_ shifts with respect to model values is 0.80. We demonstrated that a finite-size correction or much enlarged simulation box size can remove a systematic error of the calculated p*K*_a_’s and improve agreement with experiment. Importantly, the simulations captured the relevant biology in several challenging cases, e.g., the titration order of the catalytic dyad Glu35/Asp52 in HEWL and the coupled residues Asp19/Asp21 in SNase, the large p*K*_a_ upshift of the deeply buried catalytic Asp26 in thioredoxin, and the large p*K*_a_ downshift of the deeply buried catalytic Cys283 in HMCK. We anticipate that PME-CpHMD offers proper pH control to improve the accuracies of MD simulations and enables mechanistic studies of proton-coupled dynamical processes that are ubiquitous in biology but remain poorly understood due to the lack of experimental tools and limitation of current MD simulations.

## 1 Introduction

Accurate and efficient molecular modeling of proton-coupled dynamic processes is important, as biological functions and material properties often depend on protonation and deprotonation. For example, many secondary active membrane transporters utilize pH gradient and proton coupling to drive the conformation transitions for function.^1^ Many enzymes have pH-dependent catalytic activities, e.g., the active site of SARS-CoV-2 main protease collapses upon protonation of a conserved histidine residue.^2,3^ Well known examples of pH-dependent materials include aminopolysaccharide chitosan which self assembles into hydrogels in response to a small increase in solution pH.^4^ The ability to model proton-coupled dynamic processes is also important for studying protein-ligand binding, as upon protein-ligand association, the protonation state of the protein and the ligand may change.^5,6^

Unlike the conventional molecular dynamics (MD) that assumes fixed protonation states, constant pH MD allows protonation states to evolve with time according to the conformational environment and a preset solution pH. Currently, perhaps the most popular constant pH approaches are based on *λ* dynamics and the hybrid Monte-Carlo (MC)/MD scheme (also known as the stochastic titration method^7^). The former^8–12^ uses continuous titration coordinates to propagate protonation states based on an extended Lagrange approach called *λ* dynamics,^13^ while the latter^7,14–16^ combines MD with periodic MC sampling of discrete protonated and deprotonated states. Hereafter, we will refer to the former as the continuous and the latter as the discrete constant pH methods. The details of these techniques can be found in the recent reviews.^17–19^ Although the first (discrete) constant pH method^7^ is based on a hybrid-solvent scheme (see later discussion), the early implementations of the constant pH methods are solely based on the implicit-solvent generalized Born (GB) models, i.e., both conformational and protonation state sampling is conducted in implicit solvent.^8,9,14^ The use of the implicit-solvent models significantly reduces system size and allows faster sampling of solute conformational states relative to simulations with explicit water models.^17^ However, for many biologically relevant systems, e.g., trans-membrane proteins (with heterogeneous dielectric environment), nucleic acids (highly charged), and protein-ligand and protein-protein bound states, implicit-solvent models are not sufficiently accurate. This has motivated the development of explicit-solvent constant pH methods, which include the hybrid-solvent scheme and the all-atom approaches.

In a hybrid-solvent constant pH scheme, the MC sampling of protonation states or *λ* dynamics propagation of titration coordinates is conducted in implicit solvent, while MD is conducted in explicit water. The first hybrid-solvent constant pH method was developed by Baptista and Soares, who combined MD in explicit solvent with MC sampling based on Poisson-Boltzmann (PB) calculations.^7^ This method was first implemented in GROMOS96^20^ and later improved and implemented^21^ in GROMACS.^22^ Following the aforementioned work and making use of the state-of-the-art GB models, the hybrid-solvent continuous^23^ and discrete^15^ constant pH methods were developed and implemented in CHARMM^24^ and Amber.^25^ Compared to the purely GB based constant pH methods, the hybrid-solvent approaches demonstrated significantly improved accuracy for conformational dynamics and consequently better agreement with the experimental p*K*_a_’s.^15,23,26,27^ Importantly, the hybrid-solvent approach allowed the investigations of pH-dependent mechanisms of a variety of systems that are (due to inaccuracy) unfeasible to model with implicit-solvent models, e.g., proteins in mixed solvent,^28^ phase transition of surfactants,^29^ polysaccharides,^4^ and lipid bilayers,^30^ proteins at the water-membrane interface,^31^ as well as transmembrane proteins^32^ and peptides inserted in the membrane.^33^ Nonetheless, a drawback is that the Hamiltonian cannot be expressed in a hybrid scheme (semigrand canonical ensemble), and thus energy conservation is not proven to hold. In terms of applications, hybrid-solvent simulations of protein-ligand complexes are challenging, as the implicit-solvent description for ligand is not very accurate and the effects of explicit water and ions which play significantly roles cannot be fully modeled.^34^

To overcome the limitations of the hybrid-solvent scheme, much effort has been made in the development of all-atom constant pH methods over the past decade. The CHARMM program^24^ contains the CPU implementations of the all-atom continuous constant pH method with generalized reaction field^35,36^ or particle-mesh Ewald (PME) electrostatics for *λ* dynamics,^11^ and the multiple site *λ* dynamics (MS*λ*D)^37^ based constant pH method.^10,38^ These methods have been validated using p*K*_a_ calculations for a number of proteins^10,11,36^ as well as RNAs.^38^ The *λ* dynamics based constant pH method was also implemented in the GROMACS program,^22^ although only the single-site titratable model was considered and performance for proteins remains to be demonstrated.^12^ The NAMD program^39^ contains an implementation of the all-atom constant pH method based on a non-equilibrium MD-MC approach,^16^ which overcomes the issue of low acceptance of MC moves due to a large energy change resulting from a sudden switch in protonation state, as in the aforementioned hybrid MC/MD constant pH approaches.^15,21^

The aforementioned all-atom continuous constant pH methods^10,11,35,38^ are promising; however, the CPU implementations limit the simulation time scale and system size that can be studied. Recently, the Brooks group developed the basic lambda dynamics engine (BLaDE) which enables GPU acceleration for MS*λ*D based alchemical free energy calculations and constant pH simulations.^40^ In this work, we report the development and validation of the GPU all-atom continuous constant pH method in the *pmemd*.*cuda* engine of Amber program (version 2020).^41^ Following the discussion of the model parameterization and validation, we present the data from the p*K*_a_ calculations of benchmark proteins, including BBL, HEWL, SNase, RNase A, BACE 1, thioredoxin, and HMCK. In addition to comparison to experimental p*K*_a_ values, we will discuss the coupled titration of catalytic residues, pH-dependent response of solvent exposure, titration of deeply buried sites. Finally, we will discuss the finite-size effects and project future directions.

## 2 Methods and Implementation

### The all-atom PME continuous constant pH MD (CpHMD) method

In contrast to the conventional MD, the continuous constant pH MD (CpHMD) method treats the protonation states of titratable sites as dynamic variables {*λα*} and propagates them simultaneously with the spatial coordinates using an extended Hamiltonian,^8,42^

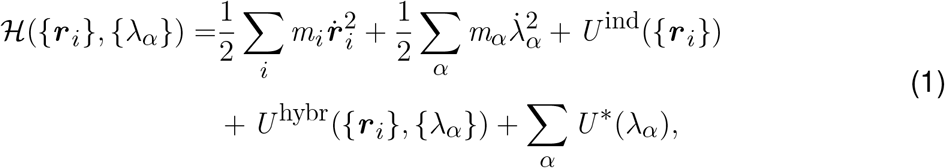

where {***r***_*i*_} and {*λ*_*α*_} refer to the spatial and titration coordinates, respectively. A deprotonated state is represented by the *λ* values close to 1 (*λ >* 0.8 in this work), whereas a protonated state is represented by the *λ* values close to 0 (*λ* ≤ 0.2 in this work). In order to impose the boundaries 0 and 1 for *λ*, we express it as^11,23,42^

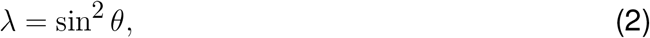

where the *θ* variable is allowed to assume any real value, as with the spatial coordinates. Therefore, *θ* is the actual coordinate in the integrator. However, for the convenience of discussion, we will write all equations in terms of *λ*.

The two first terms in the Hamiltonian (Eq. 1) describe the kinetic energies of the real atoms and *λ* particles. The *λ* particles are assigned a fictitious mass, which is similar to a heavy atom (10 amu). *U* ^ind^ represent the *λ*-independent bonded energies (see later discussion) and non-bonded energies. For the all-atom CpHMD method, *U* ^hybr^ is a sum of the *λ*-dependent Lennard Jones and electrostatic energies.^11^ The last term *U* ^*∗*^ in the Hamiltonian (Eq. 1) represents three biasing potentials that are only dependent on *λ*,

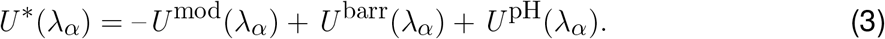

*U* ^mod^ represents the potential of mean force (PMF) for titrating a model compound or peptide in solution, which can be obtained from the traditional free energy simulations such as thermodynamic integration (TI). *U* ^barr^ is a quadratic barrier potential centered in the middle of the *λ* coordinate to prolong the residence times of the end states (*λ* close to 0 or 1):

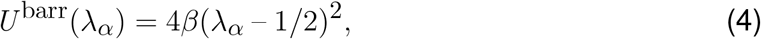

where *β* is a parameter affecting the barrier height. In the current implementation, it is set to 2.0 kcal/mol for all types of residues, similar to our previous work.^11^ *U* ^pH^ represents the free energy added to the deprotonation reaction due to a change in solution (infinite proton bath) pH

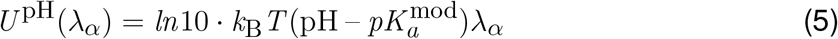

where *k*_B_ is the Boltzmann constant, *T* is the system temperature, and p*K* a^mod^ is reference p*K*_a_ of the model compound or peptide.

When *λ* = 0, the proton is present and fully interacts with its environment, and when *λ* = 1, it is treated as a ghost particle without non-bonded interactions with its environment. The partial charges on the titratable residue are linearly scaled between the protonated and deprotonated states, as in the original CpHMD framework.^8,42^ This differs from the multi-site *λ* dynamics^37^ dynamics based MS*λ*D CpHMD method,^10,38^ which scales potential energies. Formally, the *λ*-dependent Lennard-Jones interaction energy between a titratable hydrogen *i* and another non-titratable atom *j* is given by

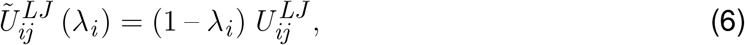

where *λ*_*i*_ is the titration variable associated with the titratable hydrogen, and 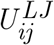 is the Lennard-Jones interaction energy between atoms *i* and *j* when the hydrogen is present. Similarly, the Lennard-Jones interaction energy between two titratable hydrogens is given by

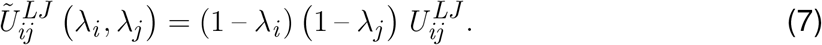

The charge of atom *j* in the titratable residue *α* is given by

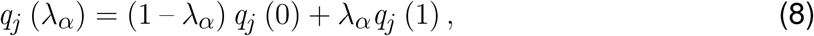

where *q*_*j*_ (0) is the charge appropriate to the protonated form of the residue, and *q*_*j*_ (1) is the charge appropriate to the deprotonated form.

The implementation presented in this paper uses fully explicit water molecules and treats the nonbonded electrostatic interaction energy between atoms with particle-mesh Ewald (PME) electrostatics. Because the *λ* values are treated as dynamic coordinates of the system, the derivatives of the energy with respect to the *λ* values are required. In the *pmemd* implementation^25^ of PME electrostatics, this interaction energy is separated into several terms,

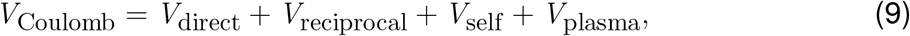

where

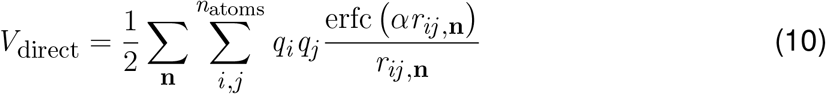

is the short-range component of the electrostatic energy, where **n** enumerates the copies of each atom from the neighboring periodic cells. This summation is performed only over those atom pairs *i, j* for which *r*_*ij*_ falls within a small cutoff distance.

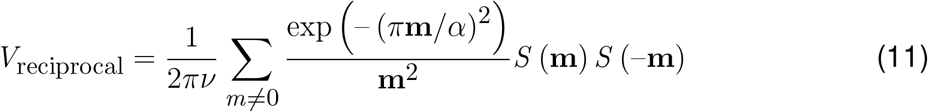

is the reciprocal space energy, where *ν* is the volume of the unit cell, **m** is reciprocal lattice vector, and *S* (**m**) is the structure factor,

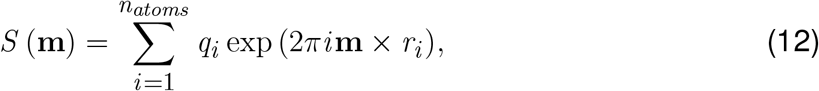

which can be approximated by

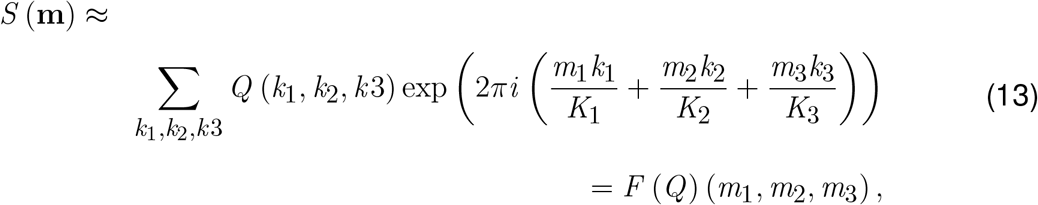

where *Q* (*k*_1_, *k*_2_, *k* 3) is a matrix constructed by interpolating the charge distribution in the simulation cell to a grid with the same dimensions *k*_1_, *k*_2_, *k*_3_, and *F* (*Q*) (*m*_1_, *m*_2_, *m*_3_) is the fast Fourier transform of the *Q* matrix.

The *V*_reciprocal_ can then be written as

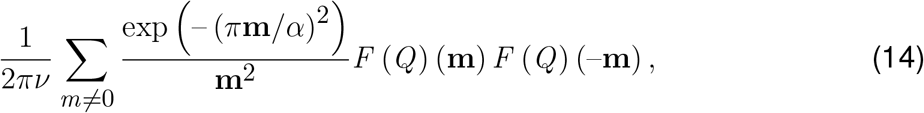

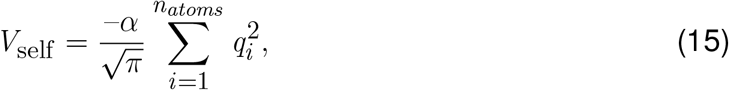

is a term that removes the self-interaction energies contained in *V*_reciprocal_, and

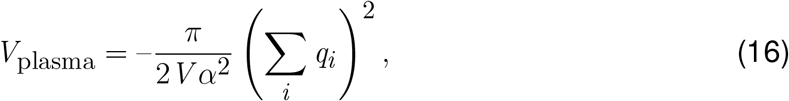

where *V* is the volume of the unit cell is a term that counterbalances any net charge on the system.

### Implementation in the *pmemd*.*cuda* engine

As in our previous CPU implementation of the PME-CpHMD method in CHARMM,^11^ the derivatives of 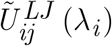 with respect to the titration variables can be derived from Eqs. 6 and 7, and computing them requires changes to be made to the Lennard-Jones forces between titratable atoms. In the present implementation, these modifications were made by making appropriate changes to the direct-force CUDA kernel in *pmemd*.*cuda* where the Lennard-Jones forces are computed. This kernel was also modified to compute the Lennard-Jones contributions to the forces on the *λ* titration variables. The electrostatic spatial forces on the atoms can be made to depend on the *λ* values by using the normal force calculations with the charges given in Eq. 8 according to the instantaneous values of the *λ* titration variables. Since *V*_self_ and *V*_plasma_ are independent of the spatial coordinates of the atoms, they are not computed during standard MD runs in *pmemd*.*cuda*. However, because these energies do depend on the *λ* titration coordinates, their derivatives with respect to the titration coordinates are required in CpHMD. The calculation of these derivatives was added to the kernel that interpolates the *λ*-dependent atomic partial charges, which was previously implemented by us for the generalized Born based CpHMD method.^43^ This kernel otherwise required minimal changes for the present implementation. The derivatives of *V*^direct^ with respect to *λ* are computed through appropriate changes to the direct-force kernel in *pmemd*.*cuda*. The derivatives of *V*_reciprocal_ with respect to *λ* are computed with a new kernel that computes the derivatives given by Eq. 14 using the same method as outlined in our previous CPU implementation in CHARMM.^11^

### Modification of the force field parameters

The current constant pH methods are based on single topology, i.e., titration is represented by switching on/off the charge and Lenard-Jones interactions of the dummy hydrogen as well as by transforming between the protonated and deprotonated forms of sidechain partial charges.^19^ The latter is straightforward to implement for the CHARMM force fields,^44^ as the backbone partial charges are independent of the side chain. This is however not the case for the Amber force fields,^45,46^ in which the backbone charges are dependent on the side chain protonation state. Due to the 1-4 interactions between the backbone and adjacent sidechain, this dependency makes it impossible to use a single reference scheme, i.e., one model for one type of sidechain. To circumvent this problem, we adopted the scheme used in the discrete constant pH implementation in Amber^14^ by fixing the backbone charges to the values of one protonation state (charged Asp/Glu and neutral His in our implementation) and absorbing the residual change in charge (ranging from 0.10 to 0.14 *e* for Asp, Glu and His) onto the C*β* atom. Such a scheme is not ideal and might introduce potential artifacts to conformational dynamics; thus, we only adopted it for titration dynamics. For conformational dynamics, the partial charges are unmodified and the charge interpolation between the protonated and deprotonated states is made to both backbone and sidechain atoms. Here we note that in our approach the conformational dynamics and titration dynamics are treated on an equal footing, with both sets of coordinates propagating together. We don’t separate these into separate phases of the simulation. Simply, we use different charge sets for the forces on the titration and spatial coordinates. By doing so, the conformational dynamics from the optimized force field is preserved.

Another compromise and approximation we made is in the bonded terms, which are not scaled between two protonation states as in the early CpHMD implementations.^8,42^ For Asp and Glu, the bonded parameters for the protonated and deprotonated forms are different in both CHARMM and Amber force fields.^44,46^ The parameters of the deprotonated forms (which are most common at physiological pH) were used except for those related to the dummy hydrogens, which were taken from the protonated forms. For His, the bonded parameters for the protonated and deprotonated forms are the same in the Amber ff14SB^46^ and CHARMM c22^44^ force fields. Including the bonded terms in the calculation would require significant modifications to the code, and as such is deferred to the future work. However, from a large number of application studies we have conducted, no artifacts have been observed, which may be due to our choice of adopting the dominant form (e.g., charged Asp/Glu). Therefore, we believe that the improvement with adding the bonded term perturbation may be minimal.

Finally, the use of two dummy hydrogens for Asp/Glu introduces an issue, namely, once an uncharged (ghost) dummy hydrogen rotates to the *anti* configuration, it loses the ability to titrate. This is because a ghost proton in the *anti* position is unfavorable to protonate, and due to zero force it is unlikely to move until it is protonated, as noted in the early developments of constant pH methods.^8,14^ Following our previous work,^42^ the rotation barrier of the C-O bond in the carboxylate group of Asp/Glu was increased to 6 kcal/mol to keep the dummy hydrogens in the *syn* configuration. This is a limitation, as the *anti* configuration might become favorable in some protein, although it is unfavorable in the peptide.^47^ One solution is to include both *anti*- and *syn*- positions for each oxygen as implemented in the discrete constant pH methods in Amber.^14,15^ This solution however is difficult to implement for CpHMD methods, as it would add additional variables which makes the analytic form of the model PMF impossible to derive (Eq. 18). To complicate the case, experimental evidence of *syn* vs. *anti* configuration is lacking. This is a topic that warrants future investigation.

### Potential of mean force functions for model titration

The linear response theory states that the charging free energy of an ion in polar solvent is quadratic in the charge perturbation.^48^ Thus, the PMF for protonation/deprotonation of a single-site titratable group (e.g., Cys and Lys) in explicit solvent can be approximated as a quadratic function in terms of *λ*.^11,36^

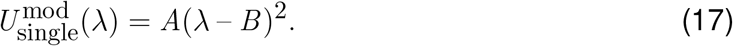

Following our previous work,^42^ for residues with two titratable sites such as carboxylic acids and histidines an additional variable *x* is introduced to represent the tautomer states. The underlying variable *θ*_*x*_ which is defined in analogy to *θ* (Eq. 2) is dynamically propagated on the same footing as *θ*. For carboxylic acids Asp and Glu which have two equivalent protonation sites (carboxyl oxygens), the model PMF function can be written as^9 42,49^

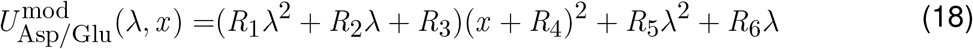

where *R*_1_, …, *R*_6_ are parameters that can be determined by one-dimensional fitting of the corresponding mean forces (*∂U* /*∂θ*)|_*θx*_ and (*∂U* /*∂θ*_*x*_)|_*θ*_ calculated using thermodynamic integration (TI) at different combinations of *θ* and *θ*_*x*_ values. The detailed derivation and protocol are given in Ref.^9,49^

The model PMF function for His titration can be written as^9^

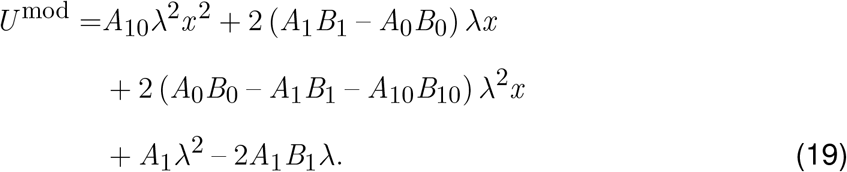

The parameters in Eq. 19 are those in the one-dimensional PMF functions, where either

*λ* or *x* is fixed at one of the end points (1 or 0).^9^

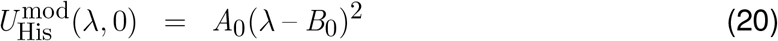

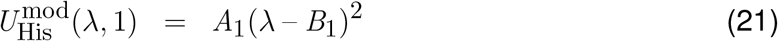

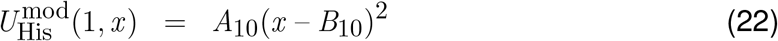

Detailed protocols for obtaining the parameters are given in Ref.^9,49^

### Finite-size corrections to the calculated p*K*_a_ values for proteins

In our previous work,^11^ we proposed a correction for the p*K*_a_’s calculated from the all-atom PME constant pH simulations under periodic boundary conditions. According to the analysis of Hü nenberger and colleagues, the finite-size errors for the ligand charging free energies arise from four physical effects, among which the discrete solvent effect dominates when the protein’s net charge is neutralized by counter-ions.^50^ The discrete solvent effect arises from a homogeneous, constant potential that is applied to offset the potential generated by isotropically tumbling solvent molecules so that the average potential over the simulation box is zero.^50^ This “offset” potential is positive for typical three-site water models, and needs to be corrected when calculating ligand charging free energies.^50^ Hü nenberger and colleagues developed an analytic correction to the ligand charging free energy^50^

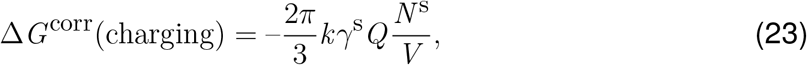

where *k* is the electrostatic constant, *γ*^s^ is the quadrupole moment trace of the solvent model relative to a van der Waals interaction site. *γ*^s^ is calculated as 0.764 *e·*Å ^2^ for TIP3P water model.^11^ *Q* is the charge (-1 for charging to -1 *e* and +1 for charging to +1 *e*), *N* ^*s*^ is the number of solvent molecules, and *V* is the simulation box volume. We note, the correction in Eq. 23 is very similar to that proposed by Roux and coworkers.^51^

Now we consider the deprotonation reactions of protein titratable residues, which refers to the charging process of an acidic sidechain, e.g., aspartic acid, Asp → Asp ^−^, or the discharging process of a basic sidechain, e.g., histidine, His^+^ → His. Based on Eq. 23 (correction for the charging free energy), we obtain the correction for the deprotonation free energy of a titratable residue in a protein in reference to a model system

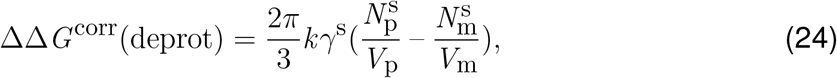

where 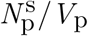 and 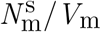 refer to the solvent number density in the protein and model systems, respectively. Note, the minus sign in Eq. 23 and *Q* are absorbed due to the fact that *Q* is -1 for acidic residues, and for basic residues Δ*G* (deprot)=-Δ*G* (charging). The corresponding p*K*_a_ correction is

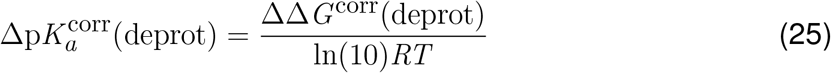

where *R* is the ideal gas constant, *T* is the temperature. Since the solvent number density is higher in the model system than in the protein system, the p*K*_a_ correction is negative for both acidic and basic sites.

## 3 Simulation Protocols

### Preparation of model peptide systems

Capped pentapeptides ACE-AAXAA-NH_2_ (X = Asp, Glu, His, Cys, or Lys) were used to parameterize and validate the model PMF functions. First, each peptide structure was generated and placed in a cubic water box using CHARMM scripts (version c38b2).^24^ The minimum distance between the heavy atoms of the peptide and the edges of the box was set as 10 Å. Next, to neutralize the system at pH 7.5, one Cl^−^ counterion was added to the Lys pentapeptide system, and one Na^+^ counterion was added to the Asp and Glu pentapeptide systems. The peptides were represented by the CHARMM c22,^44^ Amber ff14SB,^46^ or Amber ff19SB^52^ force field. Water was represented by the TIP3P water model.^53^

### Thermodynamic integration and titration simulations of the model peptides

We carried out an energy minimization in each pentapeptides system applying a force constant of 100 kcal mol^−1^Å ^−2^ to the peptide heavy atoms for 200 steps of SD followed by 300 steps of conjugate gradient method. Then, the system was heated from 100 to 300K using Langevin thermostat and a force constant of 5 kcal mol^−1^Å ^−2^ on the heavy atoms. After heating, three stages of equilibration were performed with 250 ps each, whereby the force constant was 2 and 1, and 0 kcal mol^−1^Å ^−2^. Finally, thermodynamic integration (TI) simulations were conducted for the model pentapeptides under constant NPT conditions at fixed *θ* or *θ*_*x*_ values of 0.2, 0.4, 0.6, 0.7854, 1.0, 1.2, and 1.4. Each simulation lasted 10 ns. The TI simulations gave the mean forces, ⟨*∂U* /*∂θ*|_*θx*_ ⟩ and ⟨*∂U* /*∂θ*_*x*_ |_*θ*_⟩, which were used to obtain parameters in the PMF functions (Eq. 17, 18, and 19). The detailed protocols are given in a recent tutorial.^49^

As validation of the model parameters, titration simulations were conducted for the model peptides at independent pH conditions, which were placed at 0.5-pH intervals in the range of 2–5.5 for Asp, 2.5–6 for Glu, 4.5–8 for His, 6.5–10 for Cys, and 8–11.5 for Lys model peptides. The equilibration and production runs of the peptide systems followed the same protocols as the protein simulations (see latter discussion). The production run at each pH lasted 20 ns and was repeated three times. With the CHARMM c22 force field,^44^ we also performed pH replica-exchange simulations of 10 ns/replica for Asp, His, and Lys model peptides with the same pH conditions. Additional pH replica-exchange simulations were also performed with the hydrogen mass repartition scheme^54,55^ and 4-fs timestep. The simulation length was 10 ns/replica.

### Preparation of the protein systems

For protein simulations, the following PDB files were downloaded: 1W4H (peripheral subunit-binding domain protein BBL),^56^ 2LZT (hen egg white lysozyme or HEWL),^57^ 3BDC (Staphylococcus nuclease or SNase),^58^ 7RSA (ribonucleas A or RNaseA),^59^ 1ERU (thioredoxin),^60^ and 1I0E (human muscle creatine kinase or HMCK).^61^ The coordinates were first processed using the convpdb.pl script from MMTSB Toolset^62^ to remove hetero atoms, ions, water, ligands, and hydrogen atoms. The CHARMM c22 protein force field and CHARMM modified TIP3P water model were used to represent the protein and water, respectively.^44^ The following steps were performed using the CHARMM package (c38b2).^24^ The proteins were embedded in a pre-equilibrated cubic TIP3P water box with at least 10 Å cushion between the protein heavy atoms and the edges of the box. Sodium and chloride ions were added to neutralize the systems (assuming model p*K*_a_’s and pH 7.5) and to provide a physiological (0.15 M) or experimental salt concentration (0.1 M for SNase, 0.5 M for thioredoxin, and 0.06 M for RNase A). Using the HBUILD facility, missing hydrogens were added, and a custom CHARMM script is used to add two dummy hydrogens on the carboxylate oxygens.^9^ The protein structures were energy minimized using 50 steps of steepest descent (SD) method with a harmonic force constant of 50 kcal*·* mol^−1^Å ^−2^ on the heavy atoms followed by 100 steps of adoptive basis Newton-Raphson (ABNR) method.

### Equilibration of the protein systems at independent pH conditions

The CHARMM22 topology and parameter files were converted to the Amber compatible format with the command chamber in ParmEd.^63^ With the Amber input files prepared, a last round of minimization was performed in Amber22,^25^ using 200 steps of SD followed by 300 steps of conjugate gradient method, whereby a force constant of 100 kcal mol^−1^Å ^−2^ was applied to the protein heavy atoms. Keeping the same restraint and with a time step of 1 fs, the system was then heated for 100 ps from the initial temperature of 100 K to 300 K using Langevin thermostat). Following heating, two stages of equilibration was performed. The first stage consisted of two runs of 250 ps each performed at pH 7, whereby the harmonic force constant was 100 and 10 kcal *·* mol^−1^Å ^−2^. The second stage of equilibration was performed at the individual pH conditions of the replica-exchange simulations. Here, four runs of 500 ns were performed using a time step of 2 fs. The heavy-atom force constant was gradually reduced from 10.0 to 1.0, 0.1, and 0.0 kcal mol^−1^Å ^−2^.

### Production CpHMD simulations of proteins with pH replica-exchange

For CpHMD production runs, the asynchronous pH replica exchange algorithm^64^ was employed to accelerate sampling convergence of conformational and protonation states and accelerate p*K*_a_ calculations.^23^ 2 NVIDIA GTX 2080 Ti GPU cards were used. The pH range of the protein simulations was extended at least 1 pH unit below or above the lowest or highest experimental p*K*_a_ values, and the pH spacing was 0.5 pH unit. Additional pH replica at 0.25 pH units were added in some cases to increase the probabilities of replica exchange. The exchanges between adjacent pH replicas were attempted every 2 ps (1000 MD steps). Each replica in the simulations of BBL, HEWL, SNase, thioredoxin, RNase A, and HMCK was run for 34, 40, 40, 50, 40, and 30 ns, respectively. The simulation length was sufficient to converge the p*K*_a_’s of all titratable sites (for HMCK we were only interested in Cys283). For SNase, additional simulations with larger box sizes were carried out. In these systems, the distance between the protein and edges of the water box was increased from the default 10 Å to 12, 14, and 18 Å, and the corresponding simulations lasted 20, 20, 60, and 75 ns per replica, respectively. All settings in the CpHMD are identical to our previous work.^11,43^

### Settings in the MD

Unless otherwise noted, the integration timestep in the production runs was 2 fs. Lennard Jones energies and forces were smoothly switched off over the range of 10–12 Å. For long-range electrostatics, the PME method was used with a real-space cutoff of 12 Å and grid spacing of 1 Å. Each pH replica simulations was performed under constant NPT conditions, where the pressure was maintained at 1 atm by the Berendsen barostat with a relaxation time of 0.1 ps and the temperature was maintained at 300 K by the Langevin thermostat with a collision frequency of 1.0 ps^−1^.^25^

### p*K*_a_ calculation

*λ* coordinates from the titration simulations were post-processed to calculate p*K*_a_ values. Following our previous definition of protonated and deprotonated states,^43^ *λ* ≤ 0.2 and *λ* ≥ 0.8 represent the protonated and unprotonated states, respectively, while 0.2 *< λ <* 0.8 is considered unphysical and discarded. The unprotonated fractions *S* at all simulation pH conditions were collected and the data were fit to the Hill (or generalized Henderson-Hasselbalch) equation

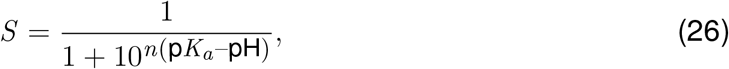

where p*K* ^a^ and *n* are the fitting parameters, and *S* is defined as *S* = *N*_unprot_/*N*_unprot_ + *N*_prot_, where *N*_unprot_ and *N*_prot_ refer to the number of *λ* values representing the unprotonated and protonated states, respectively.

For two residues experiencing linked titration, the average number of protons bound to the two residues (*(P)*) are calculated at all simulation pH, and fit to the following coupled titration model to determine the macroscopic stepwise p*K*_a_’s,^35,65^

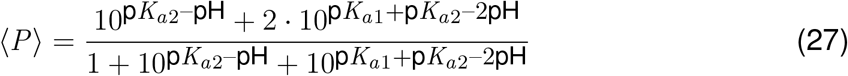

where p*K*_*a*1_ and p*K*_*a*2_ are the two stepwise p*K*_a_’s.

### Finite-size corrections

A finite-size correction (Eq. 25) was applied to the calculated p*K*_a_’s. For the p*K*_a_’s in Table 2 (i.e., a minimum of 10 Å distance between the protein and the edge of the water box), the corrections are: BBL (Asp: -0.33, Glu: -0.39, His: -0.30); HEWL (Asp: -0.63, Glu: -0.70, His: -0.61); SNase (Asp: -0.70, Glu: -0.77, His: -0.67); thioredoxin (Asp: -0.96, Glu: -1.02, His: -0.93); RnaseA (Asp: -0.66, Glu: -0.72, His: -0.63); creatine kinase (Cys: -0.64). Corrections for the simulations with larger box sizes (Table 3) are given in Table S1. At a first glance, it may seem odd that these corrections differ by residue type. This is because the corrections for the model p*K*_a_’s are different. In the future, these differences can be eliminated by using larger solvent boxes for the model simulations. Additionally, the reference p*K*_a_’s can be adjusted to account for the p*K*_a_ corrections which can be calculated at the simulation set up by using lattice parameters.^11^

## 4 Results and Discussion

### 4.1 Model parameterization and validation

#### Parameterization of the model potential of mean functions for titrating model pentapeptides

First, TI simulations of model pentapeptides CH_3_CO-Ala-Ala-X-Ala-Ala-CONH_2_ (X=Asp, Glu, His, Cys or Lys) were performed to obtain the mean forces, ⟨*∂U* /*∂θ*⟨ |_*θx*_ and/or ⟨*∂U* /*∂θ*_*x*_ ⟩ |_*θ*_), which were then fit to the analytic functions (derivatives of Eqs.17, 18, and 19 expressed in *θ*) to obtain the parameters. The fitting was generally very good (see an example fitting of His in Fig. 1), suggesting that the linear response theory holds, consistent with the results of both the GRF-based and PME-CpHMD in CHARMM.^11,36^ Integration of the mean forces followed by coordinate transformation gives the PMF as a function of *λ* (see examples in Ref^36^). We note, the parameters are more accurate when they are derived from fitting the mean forces (as in our early work^23,42^ rather than the PMF (as in the PME-CpHMD implementation in CHARMM^11^).

**Figure 1:**
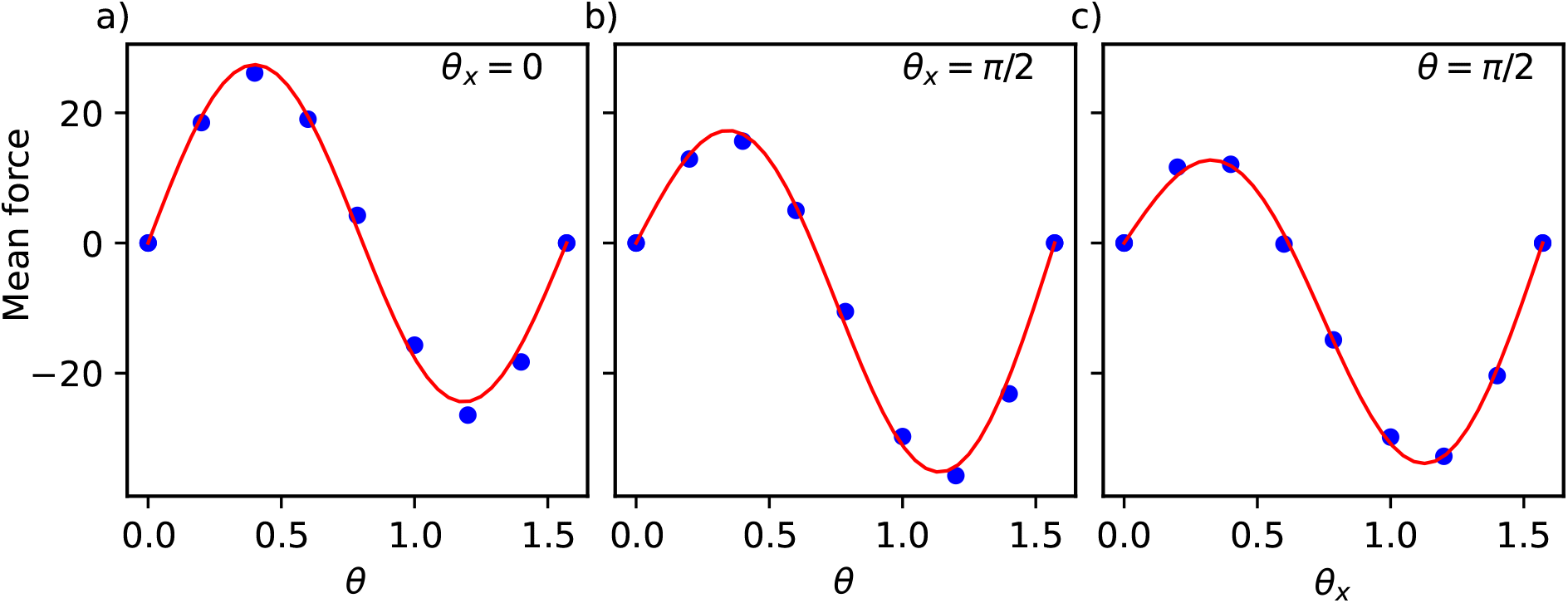
Nonlinear fitting of the mean forces to obtain the model PMF parameters for His titration. a) and b) Fitting ⟨*∂U* /*∂θ⟩* at *θ*_*x*_ = 0 (a) or at *θ*_*x*_ = *π*/2 (b) to 2*A*_0_(sin^2^*θ* – *B*_0_)sin2*θ* gives *A*_0_ and *B*_0_ (a) or *A*_1_ and *B*_1_ (b), respectively. c) Fitting ⟨*∂U* /*∂θ*_*x*_ ⟩ at *θ* = *π*/2 to 2*A*_10_(sin^2^*θ*_*x*_ – *B*_10_)sin2*θ*_*x*_ gives *A*_10_ and *B*_10_. The fitting equations are derivatives of Eqs. 20, 21, and 22. The red curves are the best fits.

Table S2 gives the parameters in the model PMF functions of Asp, Glu, and His (Eq. 18 and 19) for the CHARMM c22,^44^ Amber ff14sb,^46^ and ff19sb^52^ force fields. The model PMF parameters for Cys titration were also obtained for the CHARMM c22^44^ and Amber ff14sb^46^ force fields (Table S2). In the rest of the paper, we focused on the CHARMM c22 force field^66^ to facilitate comparisons with our previous PME-CpHMD^11^ and the Brooks’ lab’s MS*λ*D CpHMD implementations^10,37^ in CHARMM.^24^ A force field comparison study will be conducted in the near future.

#### Independent pH and replica-exchange simulations of model pentapeptides

The PMF function obtained from the TI simulations describes the free energy change along *λ*, and the difference between the two end points (*λ*=1 and 0) gives the deprotonation free energy. If the latter is reproduced by the CpHMD simulation, *λ* should sample two end (protonated and deprotonated) states with equal probabilities when pH is set to the reference p*K*_a_ value. In other words, the p*K*_a_ calculated from the titration simulation should be the same as the reference p*K*_a_. To test it, we carried out titration simulations of model pentapeptides at 8 independent pH conditions. Three replica runs of 20 ns each were performed at each pH. The unprotonated fractions at all pH conditions are converged (see time series analysis in Fig. S1). Fitting the unprotonated fractions to the Henderson-Hasselbalch equation (Eq. 26) gave the p*K*_a_’s of 3.4*±*0.04, 4.2*±*0.02, 6.5*±* 0.12, 8.4*±*0.03, and 10.3*±*0.01 for Asp, Glu, His, Cys, and Lys, respectively (Fig. 2). Fitting to the generalized Henderson-Hasselbalch equation gave identical p*K*_a_’s and error estimates, but revealed a small underestimation of the Hill coefficient for all but Cys model peptides. Except for Asp, the calculated p*K*_a_’s are within 0.1 unit of the target experimental values (Table 1). His has two titratable nitrogens and hence three protonation states: the doubly protonated Hip (charge +1) and two neutral tautomers, with a proton on either N*δ* or N ϵ, These tautomer are respectively named Hid and Hid in Amber^25^ or HSD and HSE in CHARMM.^24^ The calculated p*K*_a_’s of N*δ* (Hip ⇌ Hie) and N (Hip ⇌ Hid) are 7.0*±*0.11 and 6.7*±*0.12, respectively. These values are also within 0.1 units from the values estimated by Tanokura based on NMR data of a model compound.^67^ The titration of Asp and His is nosier than Glu, Cys, and Lys, as evident from the larger uncertainties of the unprotonated fractions at pH conditions near the p*K*_a_ value, consistent with the larger bootstrap errors (0.09 and 0.12, see Table 1). Trajectory analysis showed that the Asp and His sidechains can form hydrogen bonds (h-bonds) with the neighboring backbone group, resulting in meta-stable states. The carboxylate group of Asp is stabilized by h-bonding with the neighboring backbone amide, which contributes to the 0.3-unit under-estimation of the target p*K*_a_ value. This behavior was previously observed in both the GB and PME-CpHMD simulations in CHARMM.^9,11^

**Figure 2:**
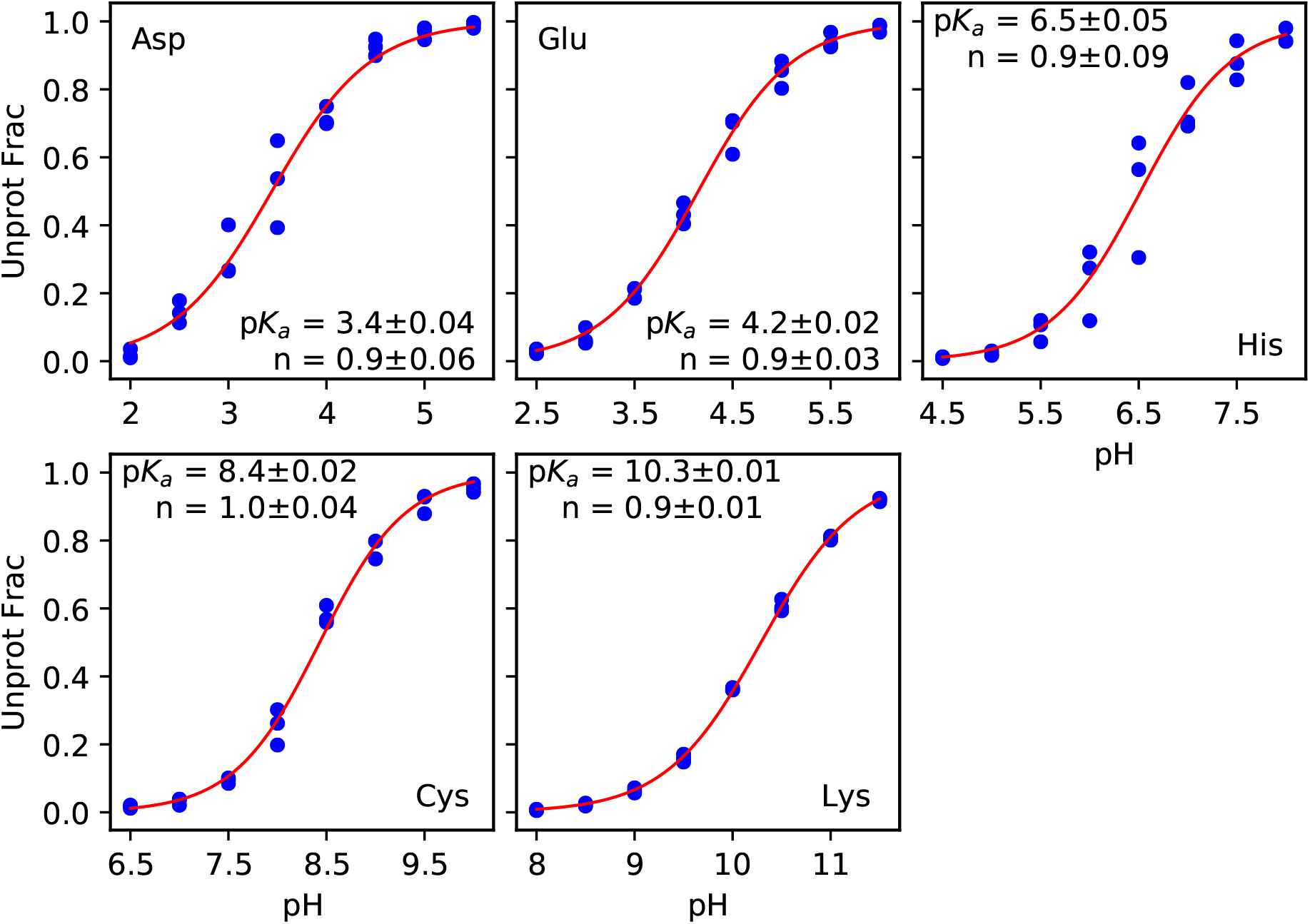
Simulated titration plots of model peptides ACEAAXAANH_2_ (X=Asp, Glu, His, Cys, and Lys) at independent pH conditions. Top panel: unprotonated fractions of Asp, Glu, and His at different pH. Bottom panel: unprotonated fractions of Cys and Lys at different pH. At each pH, three simulation runs were performed starting from different initial velocity seeds. The p*K*_a_, Hill coefficient (*n*), and fitting error are given. The boot strap errors are given in Table 2. The fitting was performed on all data points using the generalized Henderson-Hasselbalch equation. Performing the fits against the Henderson-Hasselbalch equation yields identical p*K*_a_ values and error estimates.

**Table 1:** Calculated and target experimental p*K*_a_ values of model pentapeptides

|  | Calc (IN)<sup>a</sup> | Calc (REX)<sup>b</sup> | Calc (HMR)<sup>c</sup> | Expt<sup>d</sup> |
| --- | --- | --- | --- | --- |
| Asp | 3.4±0.09 | 3.3±0.10 | 3.6±0.07 | 3.7 |
| Glu | 4.2±0.04 | | 4.3±0.02 | 4.2 |
| His | 6.5±0.12 | 6.5±0.02 | 6.3±0.03 | 6.5 |
| Hie<sup>e</sup> | 7.0±0.11 | 7.1±0.02 | 6.9±0.01 | 7.0 |
| Hid<sup>f</sup> | 6.7±0.12 | 6.7±0.03 | 6.4±0.03 | 6.6 |
| Cys | 8.4±0.03 | | 8.6±0.01 | 8.5 |
| Lys | 10.3±0.01 | 10.3±0.01 | 10.0±0.01 | 10.4 |

<sup>a</sup>Independent pH simulations, whereby each simulation was conducted for 20 ns and repeated three times. <sup>b</sup>Three sets of pH replica-exchange simulations of 10 ns/replica. <sup>c</sup>Three sets of pH replica-exchange simulations of 10 ns/replica with the HMR scheme and 4-fs timestep. All $pK_a$ 's and errors were calculated from bootstrap. <sup>d</sup>Expt refers to the NMR derived $pK_a$ 's of the model pentapeptides from Thurlkill et al.<sup>68</sup> The His tautomer $pK_a$ 's are those estimated by Tanokura based on the NMR data of a model compound.<sup>67</sup> <sup>e</sup>Hie refers to the $pK_a$ associated with Hip ⇌ Hid. <sup>f</sup>Hid refers to the $pK_a$ associated with Hip ⇌ Hie.

**Table 2:**
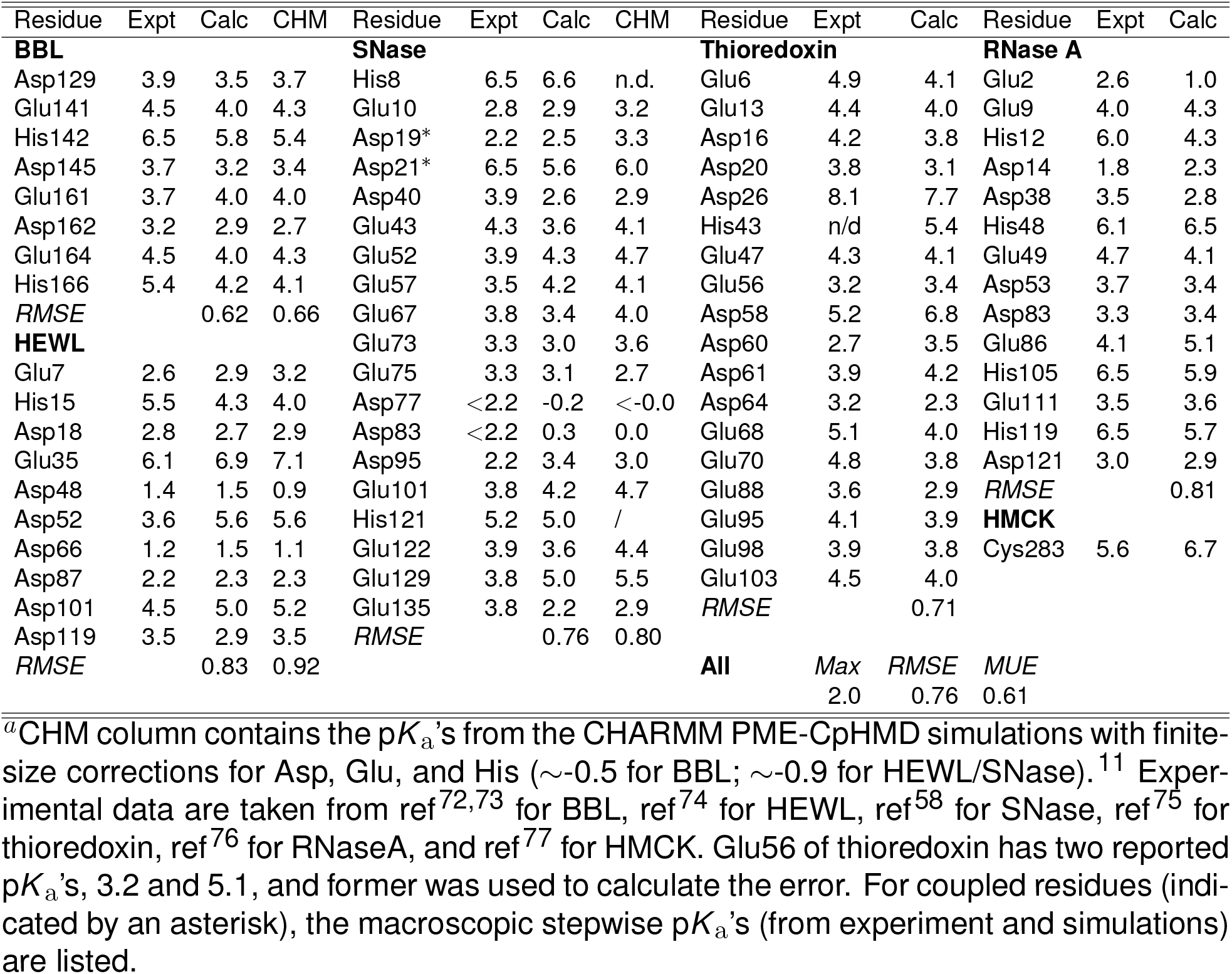
Calculated and experimental p*K*_a_’s of benchmark proteins^*a*^

To investigate if the proton-coupled conformational dynamics is adequately sampled for Asp and His in the independent pH simulations, we compared the results with those from three sets of pH replica-exchange simulations. The latter were conducted with the asynchronous pH replica-exchange scheme that was recently implemented for Amber simulations.^64^ The previous work of us^11,23,43^ and others^10,15^ demonstrated that the pH replica-exchange protocol significant accelerates protonation state and conformational sampling. Indeed, the p*K*_a_ convergence is significantly accelerated; the unprotonated fractions generally plateau after about 5 ns, compared to more than 10 ns in the independent pH simulations (Fig. S2 and S3). Interestingly, the resulting p*K*_a_’s (3.3 and 6.5) of Asp and His are very similar to those from the independent pH titration, which suggests that sampling is sufficient in the latter (Fig. S2 and S3). Note, to account for the (*∼*0.3 unit) difference between the calculated and target p*K*_a_’s of Asp pentapeptide, we changed the Asp reference p*K*_a_ (from the experimental value of 3.7 to 4.0 in the CpHMD parameter file (charmm22 pme.parm) for protein simulations.

In order to further accelerate simulations, we tested the sensitivities of p*K*_a_’s for 4-fs timestep in conjunction with the hydrogen mass repartitioning (HMR) scheme.^54,55^ Three sets of pH replica-exchange simulations of 10 ns/replica were conducted for the five model peptides with HMR/4-fs timestep. All simulations converged within 5-10 ns/replica, representing a twice speed up relative to the standard 2-fs simulations. The calculated p*K*_a_ is 3.6*±*0.07 for Asp, 4.3*±*0.02 for Glu, 6.3*±*0.03 for His, 8.6*±*0.01 for Cys, and 10.0*±*0.01 for Lys. These p*K*_a_’s deviate from the 2-fs simulations by 0.1–0.3 units. Notably, the p*K*_a_’s of the basic residues are lower, by 0.2 units for His and 0.3 units for Lys. The latter is surprising, given the rapid convergence (less than 5 ns/replica) and small random error (bootstrap error of 0.01). Trajectory analysis showed that the solvent accessible surface area (SASA) of the Lys sidechain with HMR has similar pH response, i.e., slightly decreasing with pH; however, the value for all pH conditions are higher by about 4% compared to the 2-fs simulations (data not shown). This might be related to the slightly increased diffusion constant and decreased order parameter with the 4-fs timestep, as demonstrated by a recent benchmark study.^69^ We note, evaluation of the 4-fs/HMR scheme for CpHMD simulations of proteins is not in the scope of the present work and will be conducted in the near future.

### 4.2 Titration simulations of proteins

#### Overall comparison of the calculated and experimental p*K*_a_ values

To test the accuracy of the PME-CpHMD method for modeling protonation states of proteins, we calculated the p*K*_a_’s of Asp, Glu, His, and Cys residues in BBL, HEWL, SNase, RNase A, thioredoxin, and creatine kinase (HMCK) proteins, which have been previously used to benchmark CpHMD methods.^11,43,70,71^ For a total of 67 residues, the root mean square error (RMSE) and the mean unsigned error (MUE) of the calculated p*K*_a_’s are 0.76 and 0.61, respectively, while the Pearson’s correlation coefficient *r* is 0.85 (Figure 3). A more stringent test of the p*K*_a_ prediction accuracy is to correlate the calculated and experimental p*K*_a_ shifts (Δp*K*_a_) with respect to model values, as the Δp*K*_a_ range is much smaller than the p*K*_a_ range, exposing potentially problematic cases. Encouragingly, the *r* value for Δp*K*_a_’s is 0.80, similar to the the *r* value for absolute p*K*_a_’s, suggesting that a good correlation with experimental is achieved and consistent for different residue types (see later discussion).

**Figure 3:**
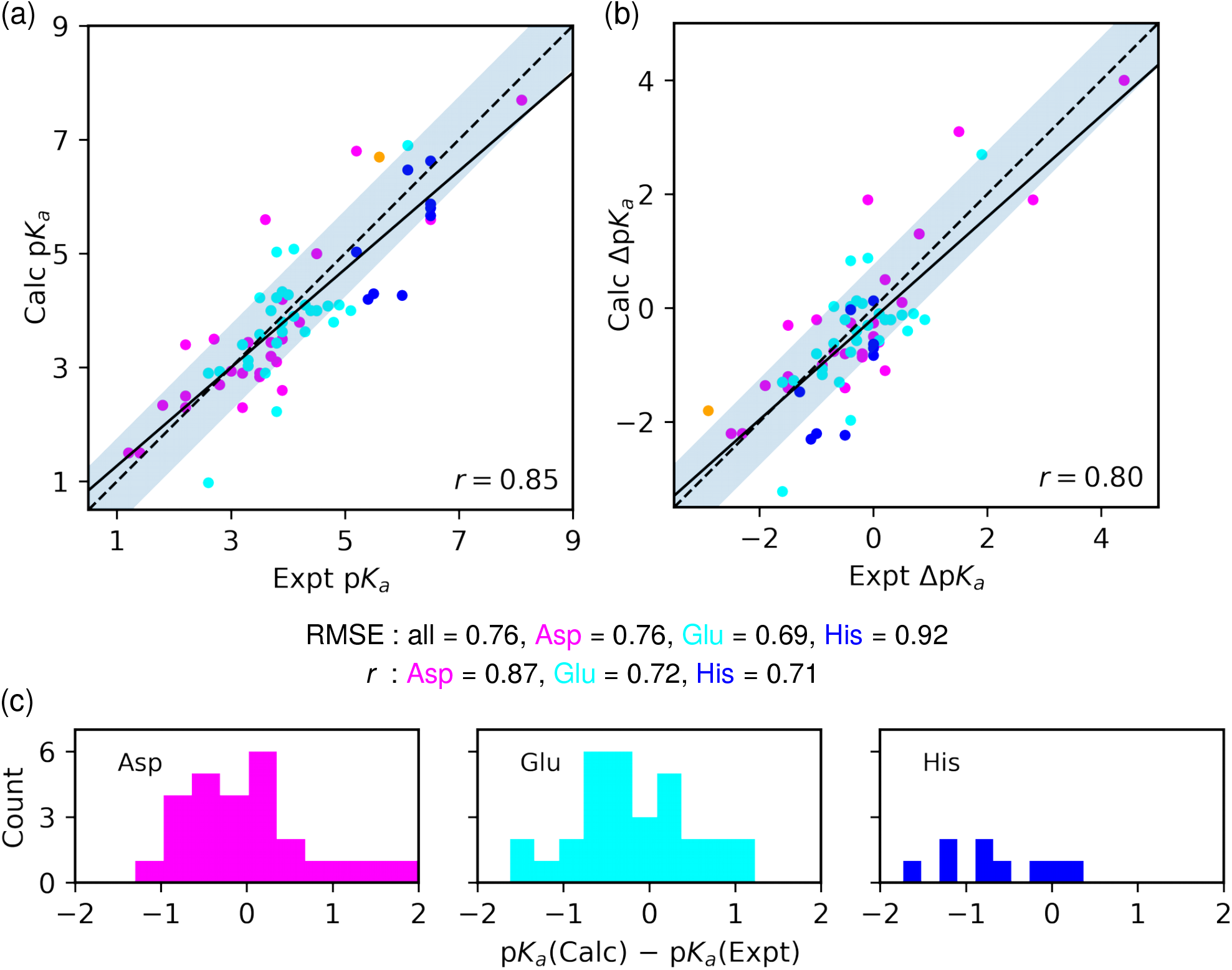
Comparison between the calculated and experimental p*K*_a_’s and p*K*_a_ shifts of the benchmark proteins. a) Calculated p*K*_a_’s vs. experimental p*K*_a_’s. b) Calculated vs. experimental p*K*_a_ shifts with respect to the experimental model peptide p*K*_a_’s (Table 1). The data for Asp, Glu, His, and Cys are shown in magenta, cyan, blue, and orange, respectively. Pearson’s correlation coefficient (*r*) and RMSE are given. The solid black lines represent the linear regression. The shaded region indicates the calculated p*K*_a_’s within the overall RMSE (0.76 units) of the experimental values. To guide the eye, the dashed diagonal line (x=y) is shown. c) Histograms of the deviations between the calculated and experimental p*K*_a_’s for Asp (left), Glu (middle), and His (right) residues.

#### Comparison of the calculated p*K*_a_’s with the all-atom CpHMD implementations in CHARMM

The p*K*_a_’s of BBL, HEWL, and SNase have been previously calculated using the all-atom PME-CpHMD implementation in CHARMM (Table S4).^11^ The *r* value of Δp*K*_a_’s (from correlation with experiment) for these proteins from the present work is 0.80, which is nearly identical to that (0.78) using the CHARMM PME-CpHMD titration.^11^ A comparison between the individual p*K*_a_’s shows that most p*K*_a_ values agree within 0.2-units (Table S4); the agreement is especially remarkable for the p*K*_a_’s of the catalytic dyad in HEWL, which differ by 0.2 units for Glu35 and are identical for Asp52.^11^ Note, the previous CHARMM PME-CpHMD simulations were run for 10 ns/replica,^11^ whereas the present Amber PME-CpHMD simulations were run until full convergence for 30–40 ns/replica. This analysis suggests that the p*K*_a_ drift is small over time and the replica-exchange CpHMD simulations offer p*K*_a_ calculations with good precision, consistent with our previous observations.^11,23^ We further compared the calculated p*K*_a_’s of BBL and HEWL, which were previously reported with the MS*λ*D method in CHARMM (Table S4).^10^ Note, the MS*λ*D simulations in Ref. used a force-based cutoff for long-range electrostatics in *λ* dynamics and therefore we did not apply a finite-size correction for the p*K* a’s.^10^ For BBL, the order of the two His p*K* a’s are in agreement between the MS*λ*D and and CpHMD results. Although the p*K* a’s from MS*λ*D are 0.6–0.8 units higher, in better agreement with experiment, the simulation length was only 5 ns/replica and no finite-size correction was applied (which downshifts the p*K*_a_’s). As to HEWL, the RMSE from MS*λ*D^10^ is nearly identical to the current work.

#### Comparison of the calculated and experimental p*K*_a_’s of different residue types

The Asp p*K*_a_’s vary the most in this dataset. Encompassing both down- and upshifted p*K*_a_’s, the experimental p*K*_a_ range for Asp is 1.2 to 8.1, similar to the calculated range of 1.5 to 7.7 (Fig. 3a, magenta). The experimental p*K*_a_’s of Glu also display both the down- and upshifts, but the range is smaller than Asp, from 2.6 to 6.1, compared to the calculated range of 1.0 to 6.9 (Fig. 3a, cyan). The overall accuracy of the p*K*_a_ calculation for Asp is slightly worse than Glu (RMSE of 0.76 and 0.69 respectively), but the *r* value for Asp (0.87) is somewhat larger than for Glu (0.72), which may be attributed to the larger p*K*_a_ range. There are only 9 experimental p*K*_a_’s for His in the current dataset, which has a range of 5.2–6.5 and do not include upshifted values (Fig. 3a, blue). The calculated His p*K*_a_ range is 4.2–6.6 (Fig. 3a, blue), and there is a trend of systematic overestimation of p*K*_a_ downshifts (Fig. 3c, right). In contrast, there is no clear trend for the p*K*_a_ errors of Asp and Glu (Fig. 3c, left and middle). The RMSE (0.92) for His p*K*_a_’s is larger than those for Asp (0.76) or Glu (0.9).

#### p*K*_a_ calculation for BBL: pH-dependent solvent exposure of His166

BBL is a miniprotein with 45 residues and 8 titratable sites. The RMSE of the calculated p*K*_a_’s is 0.62 units, with His166 showing the largest error of 1.2 units, representing an overestimation to the experimentally observed p*K*_a_ downshift (Table 2). The p*K*_a_ downshift of His166 can be attributed to solvent exclusion and lack of hydrogen bonding (h-bonding) or electrostatic interactions (Fig. 4a). As pH decreases from 6 to 4, His166 undergoes a sigmoidal transition (Fig. 4b) from the deprotonated fraction of 1 (singly protonated neutral state) to 0 (doubly protonated charged state). As expected, the fraction of the fully buried state also decreases (i.e., solvent exposure increases); however, the decrease does not appear to be sufficient, i.e., at low pH values the buried fraction doe not plateau (Fig. 4c). This analysis suggests that while the PME-CpHMD method is able to reproduce the experimental p*K*_a_ downshift by capturing the pH-induced decrease in solvent exclusion or increase in solvent exposure of His166, sampling of the exposed state at low pH may be insufficient, which contributes to the p*K*_a_ underestimation. Another potential contributor is an overestimation of the desolvation penalty by the CHARMM c22 force field.^44^ Note, our previous CHARMM PME-CpHMD simulations gave a similarly underestimated p*K*_a_ for His166 (by 1.3 units),^11^ and the MS*λ*D simulations underestimated the p*K*_a_ by 0.6 units (analysis of conformational sampling was not given).^10^

**Figure 4:**
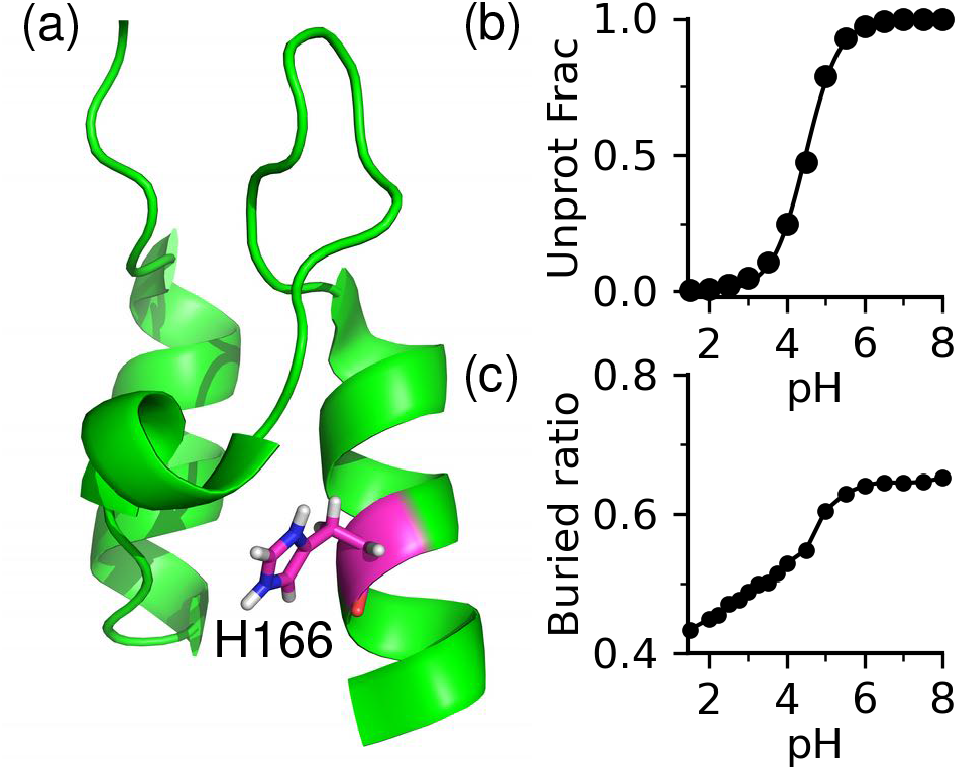
Protonation of His166 in BBL is correlated with the pH-dependent increase in solvent exposure. a) Deprotonated fraction of His166 at different pH. b) Buried ratio of His166 at different pH, defined as 1-fSASA. fSASA (fraction of solvent accessible surface area) was calculated as SASA of the sidechain atoms relative to that in the model pentapeptide.

#### p*K*_a_ calculation for HEWL: titration order of the catalytic dyad

HEWL is a small protein with 129 residues and 10 titratable sites; it is a popular test system for p*K*_a_ prediction methods due to the abundance of experimental data.^74^ The RMSE of the calculated p*K*_a_’s is 0.83 units. The two largest errors are for Asp52 and His15; the calculated p*K*_a_’s are 2 units over- and 1 unit underestimated, respectively (Table 2). Despite the overestimation, the calculated p*K*_a_ of Asp52 is 1.4 units lower than that of Glu35 (the second catalytic residue, Fig. 5), which indicates that Glu35 is a general acid and Asp52 is a general base in catalysis, in agreement with experiment (Table 2). Consistent with the CHARMM PME-CpHMD as well as the hybrid-solvent CpHMD simulations,^78^ the titration events of Glu35 and Asp52 are uncoupled, as evident from the nearly identical stepwise p*K*_a_’s (7.0 and 5.5) from fitting to the two-proton titration model (Eq. 27; figure not shown). The lack of coupled titration is due to the relatively large distance between the carboxylate sidechains (*>*6.5 Å between the nearest carboxylate oxygens at any pH). Note, the calculated dyad p*K*_a_’s are nearly the same as the values from the CHARMM PME-CpHMD simulations.^11^ The MS*λ*D method in CHARMM gave a nearly identical p*K*_a_ for Glu35, but a 1.1-unit lower p*K*_a_ for Asp52.^10^

**Figure 5:**
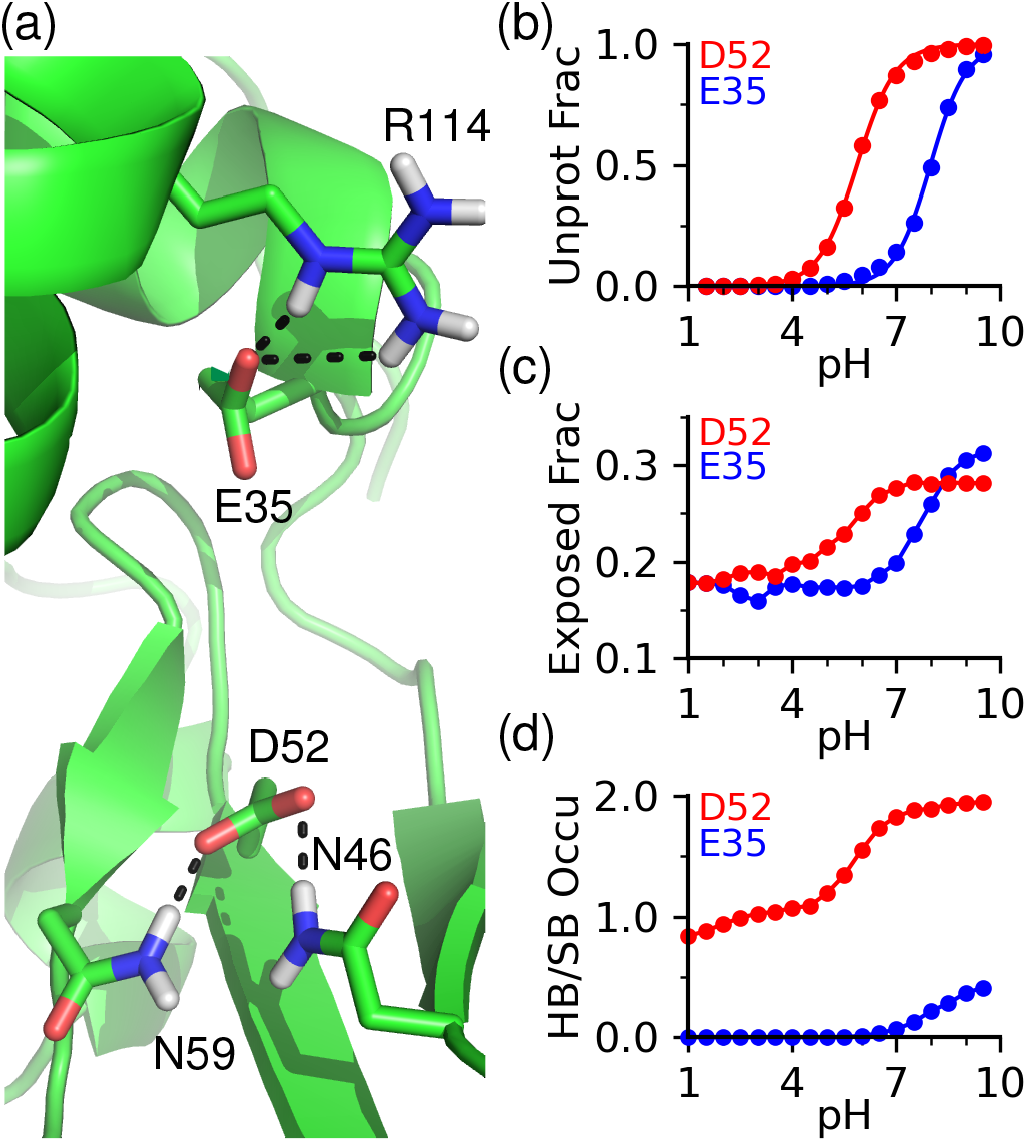
Factors influencing the p*K*_a_’s of the catalytic dyad in HEWL. (a) A representative snapshot at pH 7.5 showing the h-bonding and salt bridge environment of Glu35 and Asp52. (b) The unprotonation fractions of Glu35 and Asp52 at different pH. (c) Fractions of the sidechain solvent exposure (SASA value relative to the model pentapeptide) of Glu35 and Asp52 at different pH. (d) The h-bond and salt bridge occupancies of Glu35 and Asp52 at different pH.

To understand the p*K*_a_ order of the catalytic dayd Glu35/Asp52 and the possible factors for the p*K*_a_ overestimation of the latter, we compared the pH profiles of the solvent exposure, h-bonding and electrostatic interactions of the dyad residues with the pH-dependent titration curves (Fig. 5b–d). Both residues are partially buried. As Glu35 switches from being fully unprotonated to fully protonated in the pH range 7 to 10, the solvent exposure decreases from about 30% to just under 20% (Fig. 5b and c, blue). Asp52 has a similar behavior, except that the titration and change in solvent exposure are shifted to a lower pH range 7 to 4 (Fig. 5b and c, red). Now we turn to h-bonding and electrostatic interactions that are also physical determinants of p*K*_a_ shifts.^79^ Glu35 does not form h-bonds below pH 7.5, and above pH 7.5, occasional salt-bridge interaction with Arg114 was observed, with an occupancy less than 20% (Fig. 5d, blue). In contrast, the carboxylate group of Asp52 can accept h-bonds from the sidechains of Asn46 and Asn59, and the occupancy increases to nearly 2 with increased deprotonation of Asp52 (Fig. 5d, red). The analysis of solvent exposure and h-bond suggests that the latter is the major determinant for the lower p*K*_a_ value of Asp52 relative to Glu35, in agreement with our previous work using the hybrid-solvent as well as the PME-CpHMD in CHARMM.^78^ It is noteworthy that despite the correlation between charging of Asp52 and the increase in solvent exposure and h-bond formation, the pH profiles of solvent exposure and h-bond occupancy are more gradual than the titration curve, which may indicate that the pH-dependent conformational changes might be undersampled, contributing the overestimation of the p*K*_a_ of Asp52.

#### p*K*_a_ calculation for SNase: partially buried residues and coupled titration of Asp19 and Asp21

SNase has a large number of engineered mutants, which are popular model systems for testing p*K*_a_ prediction methods.^79^ Here we used the hyperstable, acid-resistant form of SNase Δ+PHS (hereafter referred to as SNase)^58^ which is only slightly (14 residues) larger than HEWL, but has 9 more titratable sites. The p*K*_a_’s of SNase are more challenging to predict than HEWL due to the fact that most titratable sites are partially buried.^23^ The calculated p*K*_a_’s have a RMSE of 0.76 units, similar to the RMSE of 0.80 from the CHARMM PME-CpHMD simulations.^11^ The largest error is for Glu129, for which the experimental p*K*_a_ is 0.4 units down- and the calculated p*K*_a_ is 0.8 units upshifted relative to the model value of 4.2. Curiously, simulation also fails to reproduce the direction of the experimental p*K*_a_ shifts of Glu52, Glu57, and Glu101, although the magnitude of the errors is smaller (0.4, 0.7, and 0.4 respectively). Analysis showed that all these residues are partially buried, suggesting that desolvation penalty contributes to the p*K*_a_ upshift. Based on the analysis of BBL’s His166 and HEWL’s Asp52, we hypothesized that simulation overestimates the desolvation penalty due to inadequate sampling of the solvent exposed state. To test this, we plotted the fractional SASA values vs. pH for Glu52, Glu57, Glu101, and Glu129 (Fig. S4). Deprotonation of glutamic acid is expected to induce larger solvent exposure. This is indeed the case for the more exposed residues Glu52 and Glu57 (fractional SASA about 60% at low pH), although the degree of increase is small. However, solvent exposure change with pH for the more buried residues Glu101 and Glu129, for which the fractional SASA values remain at about 40 and 20% throughout the entire pH range (Fig. S4). These data support the hypothesis that the solvent exposed state may be inadequately sampled, contributing to the desolvation related p*K*_a_ upshift for Asp and Glu.

The NMR data^58^ as well as our previous work^78^ based on the hybrid-solvent and PME-CpHMD simulations in CHARMM suggest that the titration Asp19 and Asp21 is coupled. The current simulations confirmed the strong coupling as a result of h-bond formation between the two residues (Fig. 6a). Fitting the titration data to a two-proton coupled equation (Eq. 27) gives the stepwise macroscopic p*K*_a_’s of 2.5 and 5.6 (Fig. 6b), which are in good agreement with the experimental values of 2.2 and 6.5.^58^ To assign the stepwise p*K*_a_’s to individual residues, we examine the pH-dependent probabilities of four microscopic states, doubly protonated (HH), singly protonated with proton on D19 (H–) or Asp21 (–H), and doubly deprotonated (––) states (Fig. 6c). Above pH 7, Asp19/Asp21 are in the –– state (Fig. 6c, blue). As pH decreases from 7 to 5, the probability of –– decreases, while that of the –H or H– increases. Since the –H state (cyan) is more probable than the H– state (magenta) as protonation first occurs, Asp21 receives a proton first, which means the higher p*K*_a_ should be assigned to Asp21. As pH further decreases from 5 to 2, both –H and H– states are possible; however, their combined probability decreases, while the probability of the HH state increases (Fig. 6c, red). Below pH 2, the latter state dominates. Analysis of h-bonding and electrostatic interactions (Fig. 6a) shows a network among Asp19, Asp21, Thr22, Thr41, and Arg35, consistent with the CHARMM hybrid-solvent and PME-CpHMD simulations.^43^

**Figure 6:**
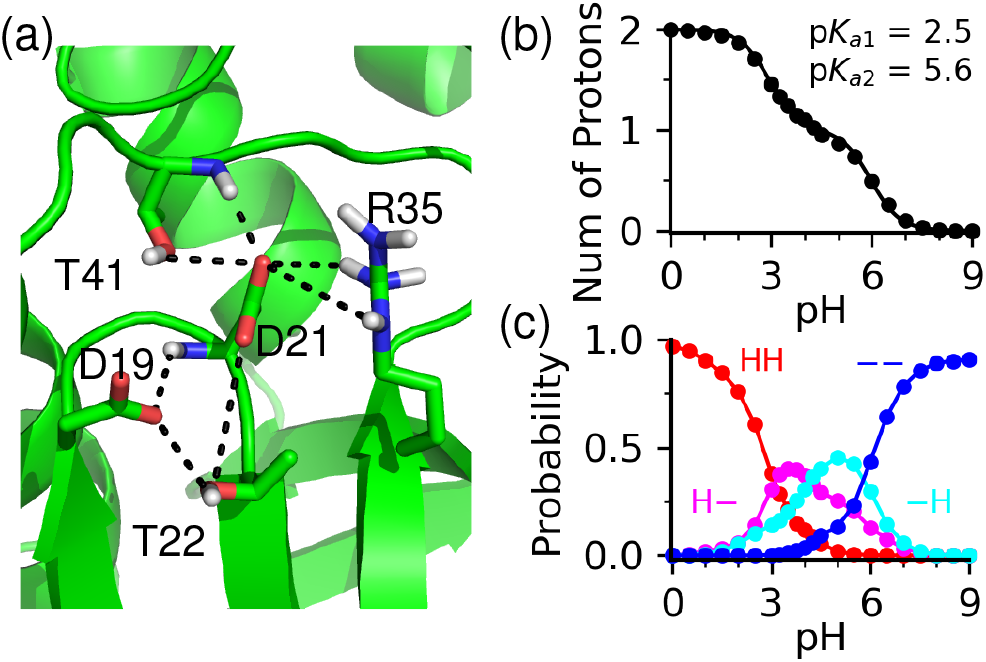
Linked titration of Asp19 and Asp21 in SNase. (a) A snapshot from the pH 4 simulation showing the h-bonding environment of Asp19 and Asp21. (b) Total number of protons of Asp19/Asp21 at different pH. The stepwise p*K*_a_’s are obtained from the best fit (black curve) to the two-proton coupled equation (Eq. 27). (c) The pH-dependent probabilities of four microscopic states: two protons (HH, red); proton on Asp19 (H–, magenta) or Asp21 (–H, cyan); zero proton (––, blue).

#### p*K*_a_ calculation for thioredoxin: the deeply buried Asp26 and Asp58

Thioredoxin has 105 residues with 18 titratable sites. The RMSE of the calculated p*K*_a_’s is 0.71. We first consider Asp26, which has one of the highest measured p*K*_a_’s of any carboxylic acids in proteins. Encouragingly, the calculated p*K*_a_ of Asp26 is 7.7, in excellent agreement with the NMR-derived value of 8.1. The large p*K*_a_ upshift of nearly 4 units can be attributed to the extremely low fraction of solvent exposure (below 7% at all pH, Fig. S5). Trajectory analysis showed that Asp26 does not form h-bonds with nearby residues. The only factor that may stabilize the deprotonated form is the salt-bridge formation with Lys39; however, the salt bridge is only formed above pH 8 and the solvent exposure is very low (*<*20% at pH 8, Fig. S5). A previous experimental study^75^ suggested that the protonated Asp26 may be stabilized by donating a h-bond to the nearby Ser28; however, in the simulation the average distance from the hydroxyl oxygen of Ser28 to the nearest carboxylate oxygen of Asp26 is about 4.9 Å at pH 7, similar to the distance of 4.7 Å in the X-ray structure (PDB 1ERU). Thus, the dry environment, along with lack of polar interactions, results in the very large p*K*_a_ upshift of Asp26.

The largest error in the calculated p*K*_a_’s of thioredoxin is for Asp58, whose direction of p*K*_a_ shift is reproduced but the magnitude is 1.6 units too large (Table 2). Analysis showed that the Asp58 is also deeply buried, with *∼*20% solvent exposure below pH 5, which explains the p*K*_a_ upshift relative to the model. However, the solvent exposure only slightly increases to *∼*30% at pH 8 before increasing steeply to over 50% at pH 10 (Fig. S6). H-bond analysis showed that the deprotonated Asp58 can accept h-bonds from the backbones of neighboring Asp60 and Asp61, which can stabilize the deprotonated state; however, the pH profile of the h-bond occupancy is irregular, showing a nearly 50% decreased occupancy in the pH range 4–8 (Fig. S6). The latter indicates a sampling issue, which explains the overestimation of the p*K*_a_ of Asp58.

#### p*K*_a_ calculation for RNase A: the deeply buried His12

RNase A has 124 residues with 14 titratable sidechains. The RMSE for the calculated p*K*_a_’s is 0.81, and the largest error is for His12 (Fig. 7a). The experimental p*K*_a_ of His12 is 0.5 units downshifted relative to the model, and the simulation overestimated the downshift by 1.7 units (Table 2). Analysis showed that His12 titrates over the pH range 3 to 6 (Fig. 7b), and the titration is correlated with two physical determinants, an increase in solvent exposure (decreased buried fraction) at lower pH (Fig. 7c) and an increase in h-bond formation at higher pH (Fig. 7d). However, unlike in the previous GB- or hybrid-solvent CpHMD simulations,^19,43^ the pH profiles of the buried fraction and the h-bond occupancy do not fully match the titration curve. Above pH 6, His12 is over 90% buried, and the decrease in the buried fraction at lower pH does not follow an expected sigmoidal curve. The buried fraction decreases by about 5% as pH decreases from 9 to 5 and remains constant between pH and 2, before further decreasing to 80% at pH 0 (Fig. 7c). A major h-bond partner is the neighboring Asn11, which can donate a h-bond from its carboxamide group to the nitrogen of His12 to stabilize its deprotonated form (Fig. 7a). As pH increases from 3 to 6, the occupancy of the h-bond increases from zero to about 60%, and it further increases to nearly 100% at pH 8. Based on the above analysis, we suggest that the h-bond formation and solvent exposure of His12 may be insufficiently sampled in the pH range 3–6. The under-sampling of the solvent exposed state (buried fraction remains unchanged between pH 2 and 4.5) is particularly evident, which may be a major factor for the overestimation of p*K*_a_ downshift of His12.

**Figure 7:**
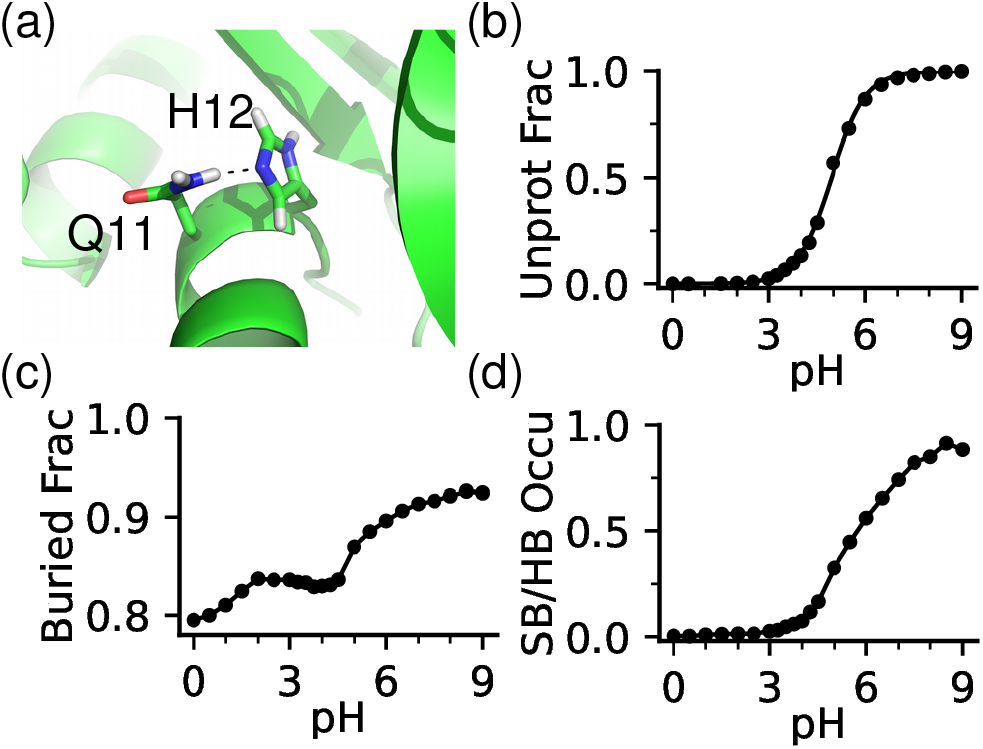
Protonation of His12 in RNaseA is correlated with the decreased solvent exclusion and hydrogen bonding. a) A zoomed-in view of the hydrogen bonding between His12 and Asn11 in RNase A. The snapshot was taken from the simulation at pH 7. b) Unprotonated fraction of His12 at different pH. c) Buried fraction of His12 at different pH. Definition of the buried fraction is given in the caption of Fig. 4. d) Occupancy of the h-bond between His12 and Asn11 at different pH.

#### p*K*_a_ calculation for HMCK: a buried active-site cysteine

To test the accuracy of Cys titration, we calculated the p*K*_a_ of Cys283 in the active site of HMCK, which has a NMR measured p*K*_a_ of 5.6,^77^ one of the lowest in the literature.^80^ Our simulations correctly reproduced the direction of the p*K*_a_ shift relative to the model; however the downshift is 1.1 units underestimated compared to the experiment (Table 2). Analysis showed that Cys283 is buried and does not have nearby cationic residues; however, once deprotonated it can accept h-bonds from the sidechains and backbones of Ser285 and Asn286 (Fig. 8a and b), consistent with the GB-based CpHMD titration simulation.^71^ Based on the structural analysis of the thioredoxin family of proteins,^81^ Roos and Messens hypothesized that hydrogen bonding rather than electrostatics plays a major role in stabilizing Cys thiolates. Our current data and recent GB-based CpHMD simulations of a large number of proteins^80,82,83^ are in support of this hypothesis.

**Figure 8:**
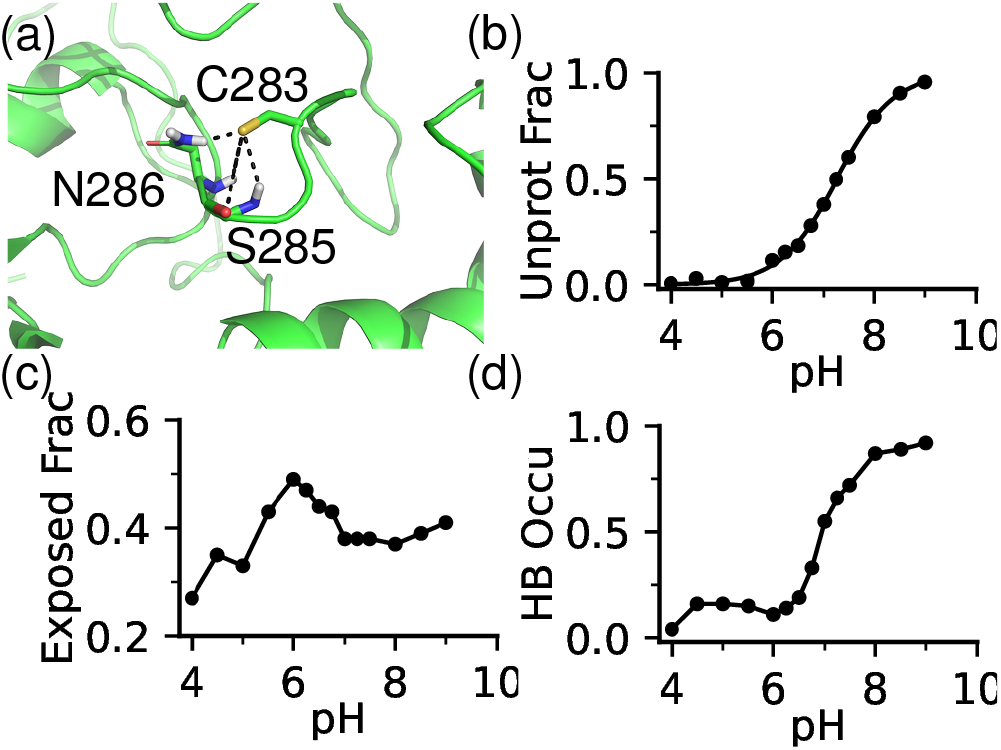
Factors influencing the p*K*_a_ of Cys283 in HMCK. (a) A zoomed-in view of the h-bond environment of Cys283 (from simulation at pH 7.5). Cys283 thiolate can form h-bonds with the backbones and sidechains of Ser285 and Asn286. (b) Unprotonated fraction of Cys283 at different pH. (c) Exposure fraction (SASA relative to that of the model pentapeptide) at different pH. d) Occupancy of the total h-bond formation of Cys283 thiolate with Ser285 and Asn286 at different pH.

As Cys283 becomes deprotonated in the pH range 6 to 9, the total h-bond occupancy increases and plateaus at 1; however, the exposed fraction does not increase and instead remains at about 40% (Fig. 8c and d). Since solvent exposure promotes the charged thiolate state and decreases the p*K*_a_, we suggest that insufficient sampling of the solvent-exposed conformations may contribute to the overestimation the p*K*_a_ of Cys283.

#### Finite-size effect and corrections

Following the work of H ü nenberger and colleagues,^50^ we previously proposed an analytical p*K*_a_ correction (Eq. 25) to correct for the effect of an offset potential introduced in PME simulations under periodic boundaries.^11^ For the current simulations, the p*K*_a_ corrections for Asp, Glu, His, and Cys are between -0.3 and -1.0 pH units (see Methods). To assess the effectiveness of the corrections and better understand the finite-size effect, we performed additional titration simulations of SNase with increased box sizes, i.e., adding more water to the simulation system. Table 3 summarizes the raw and corrected p*K*_a_’s using four different boxes, which have 10 (default), 12, 14, or 18 Å cushion space between the protein and edges of the box (minimum distance between the heavy atoms of protein and water oxygens on the box edges). The corresponding cubic box lengths are 68, 71, 76, and 84 Å, respectively.

**Table 3:** Effect of simulation box size on the calculated p*K*_a_ values of SNase^*a*^

| Residue | Expt | Raw | Corr | Raw | Corr | Raw | Corr | Raw | Corr | ΔBox |
| --- | --- | --- | --- | --- | --- | --- | --- | --- | --- | --- |
| $L_{\text{cus}}$ (Å) | | 10 | | 12 | | 14 | | 18 | | |
| $L_{\text{box}}$ (Å) | | 67.6 | | 70.6 | | 75.5 | | 83.5 | | |
| H8 | 6.5 | 7.3 | 6.6 | 7.1 | 6.5 | 7.0 | 6.5 | 7.0 | 6.7 | -0.3 |
| E10 | 2.8 | 3.7 | 2.9 | 3.5 | 2.8 | 3.5 | 2.9 | 3.1 | 2.7 | -0.6 |
| D19 | 2.2 | 3.2 | 2.5 | 3.6 | 3.0 | 3.4 | 3.0 | 3.1 | 2.8 | -0.1 |
| D21 | 6.5 | 6.3 | 5.6 | 6.5 | 5.9 | 7.1 | 6.6 | 6.4 | 6.1 | 0.1 |
| D40 | 3.9 | 3.3 | 2.6 | 2.9 | 2.3 | 2.5 | 2.0 | 2.6 | 2.3 | -0.7 |
| E43 | 4.3 | 4.4 | 3.6 | 4.4 | 3.7 | 4.3 | 3.7 | 4.1 | 3.7 | -0.3 |
| E52 | 3.9 | 5.1 | 4.3 | 5.2 | 4.5 | 4.9 | 4.3 | 4.8 | 4.4 | -0.3 |
| E57 | 3.5 | 5.0 | 4.2 | 5.1 | 4.4 | 5.2 | 4.6 | 4.4 | 4.0 | -0.6 |
| E67 | 3.8 | 4.2 | 3.4 | 4.1 | 3.4 | 4.0 | 3.5 | 3.7 | 3.3 | -0.5 |
| E73 | 3.3 | 3.8 | 3.0 | 3.6 | 3.0 | 3.3 | 2.8 | 3.1 | 2.7 | -0.7 |
| E75 | 3.3 | 3.9 | 3.1 | 3.3 | 2.6 | 3.9 | 3.3 | 3.4 | 3.0 | -0.5 |
| D77 | <2.2 | 0.5 | -0.2 | 0.6 | 0.0 | 0.5 | -0.1 | 0.4 | 0.1 | - |
| D83 | <2.2 | 1.0 | 0.3 | 2.1 | 1.5 | <0.0 | <0.0 | <0.0 | <0.0 | - |
| D95 | 2.2 | 4.1 | 3.4 | 4.1 | 3.5 | 3.8 | 3.3 | 3.2 | 2.9 | -0.9 |
| E101 | 3.8 | 5.0 | 4.2 | 4.9 | 4.2 | 4.7 | 4.2 | 4.6 | 4.2 | -0.4 |
| H121 | 5.2 | 5.7 | 5.0 | 6.1 | 5.6 | 5.7 | 5.2 | 5.8 | 5.5 | 0.1 |
| E122 | 3.9 | 4.4 | 3.6 | 4.1 | 3.4 | 4.1 | 3.5 | 4.1 | 3.7 | -0.3 |
| E129 | 3.8 | 5.8 | 5.0 | 5.8 | 5.1 | 5.5 | 5.0 | 5.4 | 5.0 | -0.4 |
| E135 | 3.8 | 3.0 | 2.2 | 3.4 | 2.7 | 3.0 | 2.5 | 3.0 | 2.6 | 0.0 |
| **RMSE** |  | 1.0 | 0.76 | 1.0 | 0.80 | 0.97 | 0.80 | 0.76 | 0.70 |  |
| **MUE** |  | 0.86 | 0.61 | 0.81 | 0.67 | 0.80 | 0.60 | 0.62 | 0.58 |  |

<sup>a</sup> Box size is represented by the solvent cushion space ( $L_{\text{cus}}$ ), i.e., minimum distance (10, 12, 14, and 18 Å) between the protein heavy atoms and edges of the water box, and the length of a cubic box ( $L_{\text{box}}$ ) converted from the average system volume. The columns Raw and Corr refer to the $pK_a$ 's before and after the finite-size corrections (see Methods). The column $\Delta$ Box refers to the raw $pK_a$ difference between the largest and smallest boxes.

We first examine the raw calculated p*K*_a_’s from simulations with different box sizes. As expected, with increasing box size the raw p*K*_a_’s decrease for all but four residues (Fig. 9a). Increasing box size also leads to better agreement with the experimental p*K*_a_’s; the RMSEs of the raw p*K*_a_’s are 1.0, 1.0, 0.97 and 0.76 for boxes with 10, 12, 14, and 18 Å cushion space, respectively (Table 3 and Fig. 9a). The MUE also decreases from 0.86 (10 Å cushion) to 0.81 (12 Å cushion), 0.80 (14 Å cushion), and 0.62 (18 Å cushion) (Table 3). Comparison of the raw p*K*_a_’s between the smallest (10 Å cushion) and largest (18 Å cushion) boxes shows that the p*K*_a_ changes due to box size increase vary (Table 3, last column). Excluding the four residues (Asp19, Asp21, H121, and Glu135) that show very little p*K*_a_ changes, the p*K*_a_’s mostly decrease by 0.3 to 0.7 units, as compared to the finite-size corrections of -0.70 to -0.8 units for the smallest box. The effect of box size is not clear for the coupled residues Asp19/Asp21, which have the raw calculated p*K*_a_’s of 3.2/6.3 with the smallest box; however, the p*K*_a_’s increase to 3.6/6.5 and 3.4/7.1 with the larger boxes (12 and 14 Å cushion space), and then decrease back to 3.1/6.4 with the largest box (18 Å cushion). Increasing box size has negligible effect on the downshifted p*K*_a_ of His121. With the increasing box sizes, its raw p*K*_a_ changes from 5.7 to 6.1, 5.7, and 5.8. Box size also shows little effect on the p*K*_a_ of Glu135, which has the raw calculated p*K*_a_’s of 3.0, 3.4, 3.0, and 3.0 with the increasing box sizes.

**Figure 9:**
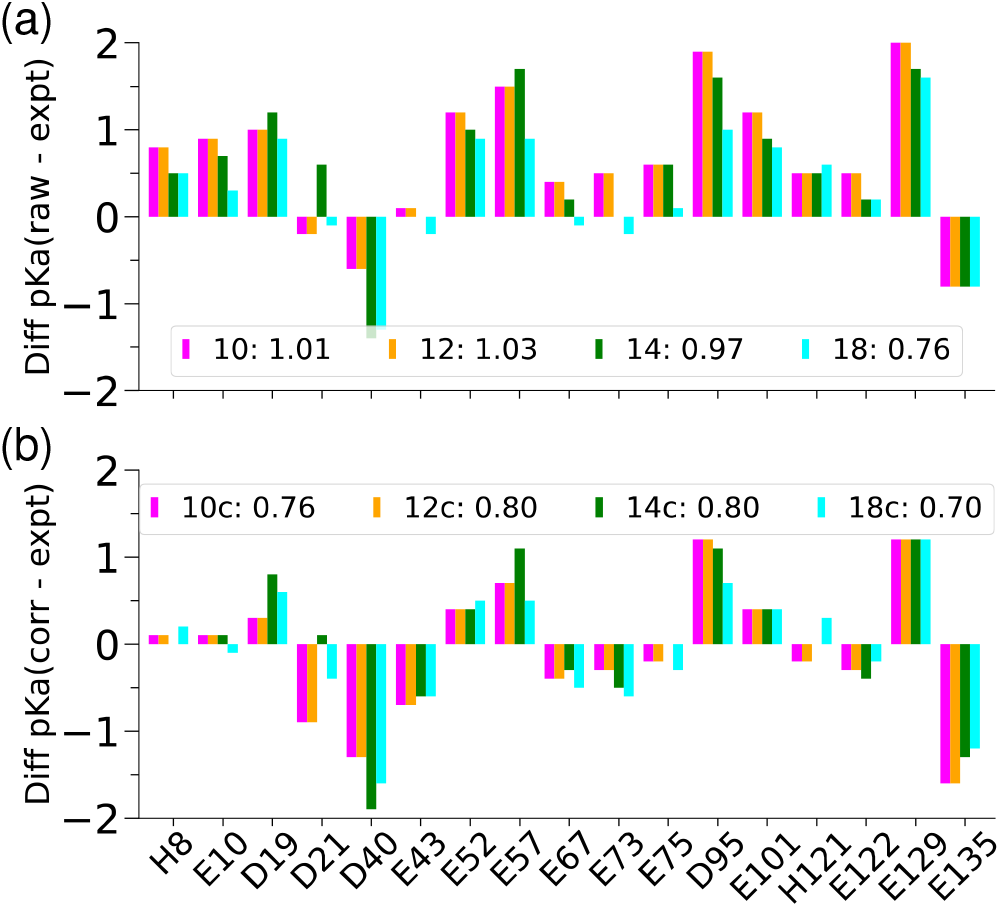
Effect of box size on the calculated p*K*_a_’s. The errors of the raw (a) and finite-size corrected (b) p*K*_a_ of SNase with different solvent cushion spaces, 10 (magenta), 12 (orange), 14 (green), and 18 Å (cyan). The corresponding RMSE values are shown next to the legends.

Now we examine the p*K*_a_ corrections for the different simulation boxes. It is apparent that application of the finite-size correction removes the systematic overshift error (Fig. 9b). As the box size increases, the solvent number density increases and therefore the magnitude of the correction decreases (Eq. 24). The magnitude of the corrections decreases by about 0.4 pH units going from the smallest to the largest box. Interestingly, this difference is roughly the same as the average difference between the raw p*K*_a_’s (of all but the aforementioned four residues) calculated with the smallest and largest box (Table 3, last column), which suggests that the finite-size correction is valid. Another interesting observation is that the increasing box size does not significantly reduce the RMSE of the finite-size corrected p*K*_a_’s. The RMSE’s are 0.76, 0.80, 0.80, and 0.70 with the increasing box sizes (Fig. 9b), which is another piece of evidence supporting the validity of the finite-size corrections.

## 5 Concluding Discussion

We presented the first implementation, parameterization, and validation of the GPU-accelerated continuous constant pH particle-mesh Ewald molecular dynamics method in Amber22 (hereafter referred to as Amber PME-CpHMD). Titration parameters for three force fields (CHARMM c22,^44^ Amber ff14SB,^46^ and ff19SB^52^) were derived and validated using model pentapeptides AAXAA, where X represents Asp, Glu, His, Cys, or Lys. To benchmark the performance and accuracy for constant pH simulations of proteins, we carried out titration simulations with the c22 force field for 6 proteins, including BBL, HEWL, SNase, RNase A, thioredoxin, and HMCK, which have NMR derived p*K*_a_ values of Asp, Glu, His, and Cys residues. The asynchronous pH replica-exchange algorithm^64^ was employed to enhance sampling of protonation and conformational states. The simulations were run for 30-50 ns per pH replica until all p*K*_a_’s were converged. The resulting RMSE and MUE with respect to the experimental p*K*_a_’s are 0.76 and 0.61, respectively, and the largest p*K*_a_ deviation is 2 units. The Pearson’s correlation coefficients for the calculated vs. experimental p*K*_a_’s and p*K*_a_ shifts are 0.85 or 0.80, respectively. Importantly, the titration simulations quantitatively reproduced the experiment p*K*_a_ orders of the catalytic dyad in HEWL and the coupled residues in SNase. Simulations also quantitatively captured one of the largest upshifted p*K*_a_’s of a deeply buried Asp in thioredoxin as well as the downshifted p*K*_a_ of an active-site Cys in HMCK.

We compared the current validation data with those based on the CHARMM^24^ CPU all-atom PME-CpHMD^11^ and MS*λ*D^10^ simulations with the same c22 force field.^44^ The Asp, Glu, and His p*K*_a_’s calculated from the CHARMM PME-CpHMD simulations of 10 ns per replica (much shorter than the present work) are in close agreement with the present work, suggesting that the p*K*_a_ drifts over prolonged simulation time are small. Comparing to the calculated p*K*_a_’s of HEWL and the two His residues in BBL based on the MS*λ*D simulations of 5-20 ns per pH replica (with a 12-Å electrostatic cutoff),^10^ the overall RMSE is similar, and the p*K*_a_ orders of the catalytic Glu35/Asp52 in HEWL and His142/His166 in BBL are consistent with the present simulations.

In agreement with the previous CHARMM PME-CpHMD simulations^11^ the present data demonstrated that the finite-size effect needs to be taken into account for the accurate calculation of titration free energies with lattice sum methods under periodic boundary conditions. Applying the p*K*_a_ correction^11^ to account for a positive offset potential due to TIP3P water in periodic boxes, a systematic p*K*_a_ upshift error in the calculated p*K*_a_’s was removed, and the overall agreement with experiment was improved. We note, in the revision stage of the current paper, the work from the Roux group^84^ was published which used a similar p*K*_a_ correction to account for the (Gavani) offset potential.^51^

To further examine the finite-size effect and the validity of the correction, the p*K*_a_’s of SNase were calculated from simulations with four different box sizes. Consistent with the negative sign of the correction, increasing box size lowers the raw p*K*_a_’s of all but four residues that do not show significant changes. The RMSE of the raw p*K*_a_’s decreases from 1.0 with the smallest box to 0.76 with the largest box; the latter is identical to the RMSE (0.76) of the corrected p*K*_a_’s obtained from the simulation with the smallest box. The quantitative validity of the correction is also supported by a good agreement between the change in the finite-size correction and the average change of the raw p*K*_a_’s going from the smallest to the largest box size. As expected, the finite-size correction decreases with increasing box size, and consistently, the reduction in RMSE due to the correction also decreases. Using the 18-Å cushion space, the correction is 0.3–0.4, and the RMSE (0.70) of the corrected p*K*_a_’s is only slightly smaller than the RMSE (0.76) of the raw p*K*_a_’s. This suggests that the box size effect may start to become negligible with this size of water box.

Although the overall box-size dependent trend of p*K*_a_’s is consistent with the positive offset potential being the dominant factor,^11^ there are exceptions. The simulations of SNase showed that box size has negligible effect on the coupled p*K*_a_’s of Asp19 and Asp21 as well as the downshifted p*K*_a_’s of His121 and Glu135. We note that the effect of the offset potential and the corresponding p*K*_a_ correction deal with an ideal situation in which a single residue titrates in a neutral background. Thus, it is possible that the correction is not valid for coupled p*K*_a_’s. However, with regards to the p*K*_a_’s of His121 and Glu135, the cause for the box size independence is difficult to speculate. An alternative approach to the finite-size p*K*_a_ correction is to enforce system charge neutrality i.e., by including titratable water as in our previous work.^11^ We tested this approach on the BBL protein; however, due to the slower convergence and small p*K*_a_ differences compared to the simulations without titratable water, studies of other proteins were not pursued. We defer a more thorough investigation of the finite-size effects to a future work.

We analyzed the pH-dependent solvent exposure and formation of hydrogen bonds as well as electrostatic interactions of catalytic residues and those that exhibit larger p*K*_a_ deviations from experiment. These analyses suggested while PME-CpHMD captures the proton-coupled conformational rearrangements, charging-induced increase of solvent exposure for buried residues is inadequate. This may be a major contributor to the p*K*_a_ errors, including the overestimated p*K*_a_ downshifts for buried His residues, e.g., His166 in BBL and His12 in RNaseA; the overestimated p*K*_a_ upshifts for buried carboxyl residues, e.g., Glu57 in SNase and Asp58 in thioredoxin; and the overestimated p*K*_a_ upshift for buried Cys, e.g., Cys283 in HMCK. Undersampling of the solvent-exposed state may also be related to the combination of c22/TIP3P force field,^44^ which slightly biased solute-solute over solute-solvent interactions.^85^ Overestimation of desolvation penalty may also be a source of error, which can be attributed to the low dielectric constant in the protein interior as a result of the lack of polarization in simulations with additive force fields.^86^ Lack of polarization in the interior of protein may also lead to overly strong salt bridges, which may explain the overestimation of the p*K*_a_ downshifts of Asp140 and Glu135 in SNase. While the use of polarizable force field for both protein and water is desirable, it may not be currently feasible due to speed. One intriguing idea worth exploring is to mix a polarizable water model such as OPC3-pol^87^ with an additive force field to improve solute-solvent interactions. The present study did not examine the potential dependence on the additive force field. The force field related topics as well as the evaluation of PME-CpHMD for model proton-coupled conformational dynamics of catalytic residues in larger proteins (e.g., BACE1^70^) will be explored in a future work.

By removing the reliance on the implicit-solvent model, the PME-CpHMD method can be applied to any system that has a force field representation. We anticipate the GPU accelerated PME-CpHMD to become a powerful tool for the investigation of a variety of proton-coupled dynamical phenomena that are poorly understood due to the current limitations in experimental and MD techniques, for example, secondary transport of ions/substrates across membrane transporter proteins and pH-dependent self-assembly of materials. Another important application of PME-CpHMD is to offer proper pH control, for example, by allowing protein and ligand to titrate while binding and unbinding,^5,6^ or allowing His residues to fluctuate among the doubly protonated and two singly protonated tautomer states, which has been shown to affect the ligand binding mechanism and kinetics.^88,89^

## Supporting Information Available

Supporting Information contains additional tables and figures.

## Acknowledgements

The authors acknowledge National Institutes of Health (R01GM098818) for funding.

