## Supplemental tables and figures for "GPU-Accelerated All-atom Particle-Mesh Ewald Continuous Constant pH Molecular Dynamics in Amber"

Table S1: Simulation box sizes and p$K_\mathrm{a}$ corrections[a]

| System | Box parameters | | | p$K_\mathrm{a}$ corrections | | | |
|---|---|---|---|---|---|---|---|
| | $N^\mathrm{s}$ | $L$ (Å) | Cushion (Å) | Asp | Glu | His | Cys |
| **Proteins** | | | | | | | |
| BBL | 6273 | 57.8 | 10 | -0.33 | -0.39 | -0.30 | |
| HEWL | 9735 | 68.6 | 10 | -0.70 | -0.77 | -0.67 | |
| SNase | 9735 | 67.6 | 10 | -0.70 | -0.77 | -0.67 | |
| SNase | 11173 | 70.6 | 12 | -0.60 | -0.66 | -0.57 | |
| SNase | 13802 | 75.5 | 14 | -0.47 | -0.54 | -0.45 | |
| SNase | 18863 | 83.5 | 18 | -0.32 | -0.39 | -0.30 | |
| Thioredoxin | 5801 | 57.3 | 10 | -0.96 | -1.02 | -0.93 | |
| RNase A | 9162 | 66.2 | 10 | -0.66 | -0.72 | -0.63 | |
| HMCK | 29059 | 97.2 | 10 | | | | -0.64 |
| **Peptides** | | | | | | | |
| Asp | 2530 | 42.4 | 10 | | | | |
| Glu | 4748 | 52.2 | 10 | | | | |
| His | 2158 | 40.2 | 10 | | | | |
| Cys | 2184 | 40.3 | 10 | | | | |

[a] $N^\mathrm{s}$ refers to the number of water in the simulation box. $L$ refers to the edge length of a cubic box converted from the average volume of the system at all pH conditions. Cushion refers to the minimal distance between the protein heavy atoms and water oxygens in the box edges. The coefficient of variation across different pH conditions is about 0.04%–0.16% for all systems.

Table S2: Parameters in the model potential of mean force functions of Asp, Glu, His, Cys, and Lys for PME-CpHMD simulations with CHARMM and Amber force fields

| Residue | R1 | R2 | R3 | R4 | R5 | R6 |
|---|---|---|---|---|---|---|
| **c22** | | | | | | |
| Asp | 0.546 | 10.135 | -10.606 | 0.500 | -74.200 | 0.105 |
| Glu | 0.800 | 9.689 | -10.453 | 0.500 | -74.169 | 0.075 |
| His | -51.698 | 0.377 | -51.784 | 0.520 | -45.356 | 0.334 |
| Cys | -81.663 | -0.004 | N/A | N/A | N/A | N/A |
| Lys | -80.323 | 0.629 | N/A | N/A | N/A | N/A |
| **ff14SB** | | | | | | |
| Asp | -0.216 | 8.855 | -8.858 | 0.500 | -46.217 | 0.224 |
| Glu | -0.676 | 11.053 | -10.533 | 0.500 | -44.808 | 0.186 |
| His | -34.781 | 0.251 | -33.445 | 0.211 | -33.506 | 0.530 |
| Cys | -77.895 | -0.040 | N/A | N/A | N/A | N/A |
| **ff19SB** | | | | | | |
| Asp | -39.809 | 88.561 | -49.157 | 0.500 | -48.830 | 0.193 |
| Glu | -37.993 | 86.876 | -49.197 | 0.500 | -48.011 | 0.160 |
| His | -34.528 | 0.214 | -33.316 | 0.180 | -33.122 | 0.531 |

Table S3: Cumulatively calculated p$K_a$'s discussed in the main text

| Time (ns) | BBL H166 | HMCK C283 | Time (ns) | HEWL E35 | HEWL D52 | SNase D19 | SNase D21 |
|---|---|---|---|---|---|---|---|
| 5 | 4.4 | 7.3 | 10 | 7.0 | 5.7 | 5.8 | 3.1 |
| 10 | 4.4 | 7.1 | 20 | 7.2 | 5.6 | 5.8 | 2.7 |
| 15 | 4.4 | 7.3 | 30 | 7.0 | 5.6 | 5.8 | 2.6 |
| 20 | 4.5 | | 40 | 6.9 | 5.6 | 5.8 | 2.6 |
| 25 | 4.5 | | 50 | | | 5.6 | 2.5 |
| 30 | 4.5 | | 60 | | | | |
| 34 | 4.5 | | 70 | | | | |

Table S4: Comparison of the p$K_a$'s calculated in this work with those from the all-atom CpHMD methods in CHARMM

| Residue | Expt | CpHMD | MS$\lambda$D | This work |
|---|---|---|---|---|
| **BBL** | | | | |
| Asp129 | 3.9 | 3.7 | / | 3.5 |
| Glu141 | 4.5 | 4.3 | / | 4.0 |
| His142 | 6.5 | 5.4 | 6.6 | 5.8 |
| Asp145 | 3.7 | 3.4 | / | 3.2 |
| Glu161 | 3.7 | 4.0 | / | 4.0 |
| Asp162 | 3.2 | 2.7 | / | 2.9 |
| Glu164 | 4.5 | 4.3 | / | 4.0 |
| His166 | 5.4 | 4.1 | 4.8 | 4.2 |
| *RMSE* | | 0.66 | | 0.62 |
| **HEWL** | | | | |
| Glu7 | 2.6 | 3.2 | 2.7 | 2.9 |
| His15 | 5.5 | 4.0 | 6.0 | 4.3 |
| Asp18 | 2.8 | 2.9 | 2.1 | 2.7 |
| Glu35 | 6.1 | 7.1 | 7.0 | 6.9 |
| Asp48 | 1.4 | 0.9 | 1.3 | 1.5 |
| Asp52 | 3.6 | 5.6 | 4.5 | 5.6 |
| Asp66 | 1.2 | 1.1 | 1.5 | 1.5 |
| Asp87 | 2.2 | 2.3 | 1.3 | 2.3 |
| Asp101 | 4.5 | 5.2 | 5.1 | 5.0 |
| Asp119 | 3.5 | 3.5 | 1.6 | 2.9 |
| *RMSE* | | 0.92 | 0.84 | 0.83 |
| **SNase** | | | | |
| His8 | 6.5 | / | | 6.6 |
| Glu10 | 2.8 | 3.2 | | 2.9 |
| Asp19 | 2.2 | 3.3 | | 2.5 |
| Asp21 | 6.5 | 6.0 | | 5.6 |
| Asp40 | 3.9 | 2.9 | | 2.6 |
| Glu43 | 4.3 | 4.1 | | 3.6 |
| Glu52 | 3.9 | 4.7 | | 4.3 |
| Glu57 | 3.5 | 4.1 | | 4.2 |
| Glu67 | 3.8 | 4.0 | | 3.4 |
| Glu73 | 3.3 | 3.6 | | 3.0 |
| Glu75 | 3.3 | 2.7 | | 3.1 |
| Asp77 | $<$2.2 | $<$-1.0 | | -0.2 |
| Asp83 | $<$2.2 | 0.0 | | 0.3 |
| Asp95 | 2.2 | 3.0 | | 3.4 |
| Glu101 | 3.8 | 4.7 | | 4.2 |
| His121 | 5.2 | / | | 5.0 |
| Glu122 | 3.9 | 4.4 | | 3.6 |
| Glu129 | 3.8 | 5.5 | | 5.0 |
| Glu135 | 3.8 | 2.9 | | 2.2 |
| *RMSE* | | 0.80 | | 0.76 |

CpHMD refers to the PME-CpHMD method in CHARMM.[S1] MS$\lambda$D refers to the $\lambda$-dynamics based constant pH method with electrostatic cutoff for $\lambda$ dynamics.[S2]

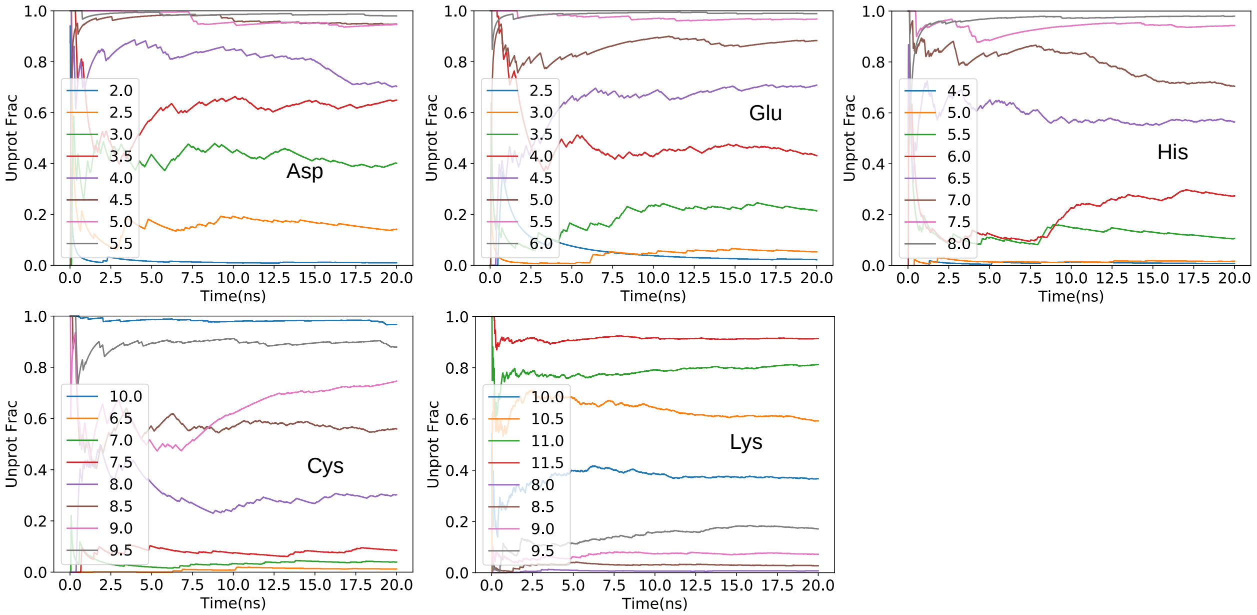


Figure S1: Time series of the cumulative unprotonated fractions of Asp, Glu, His, Cys, and Lys from the independent pH titration simulations of the model penta-peptides $CH_3COAAXAACONH_2$. Data from the replica run 1 are shown. Data from the other runs are similar.

.

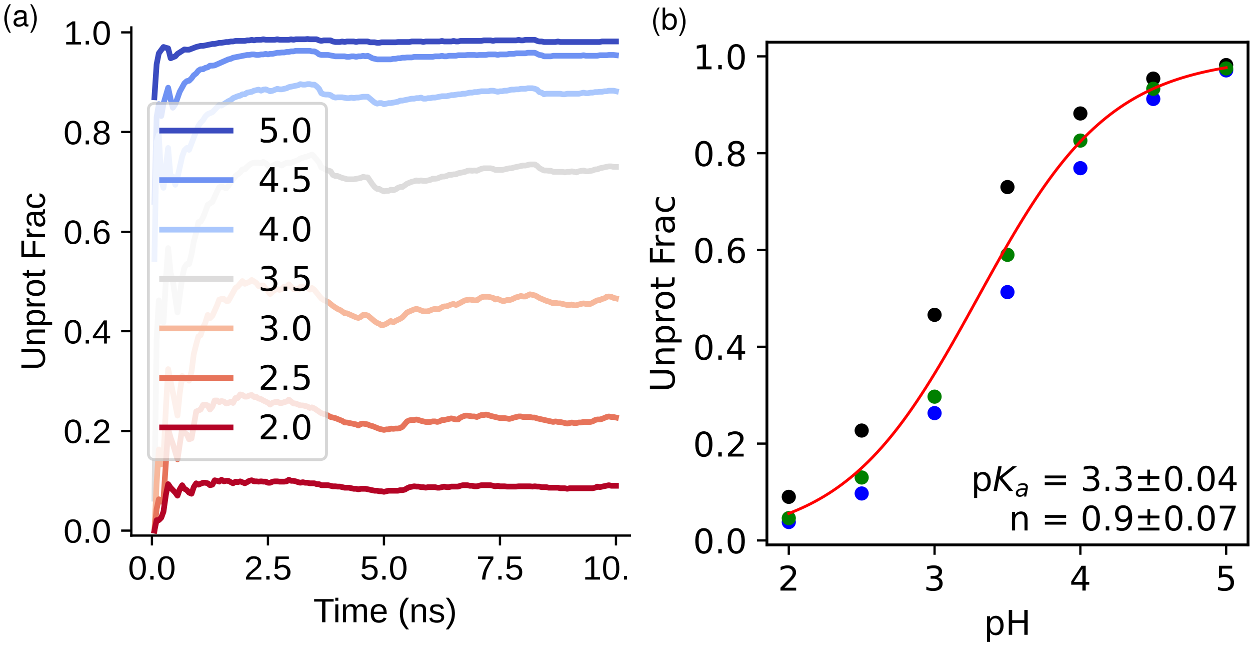


Figure S2: (a) Time series of the cumulative unprotonated fraction of Asp from the first set of three replica-exchange simulations. Legends indicate the pH conditions. (b) Unprotonated fractions of Asp at different pH. Data are from three independent sets of pH replica-exchange simulations of $CH_3COAADAACONH_2$. The best fit (red) and the p$K_\mathrm{a}$ were obtained using all data points. The p$K_\mathrm{a}$ from the bootstrap method is 3.3$\pm$0.10.

.

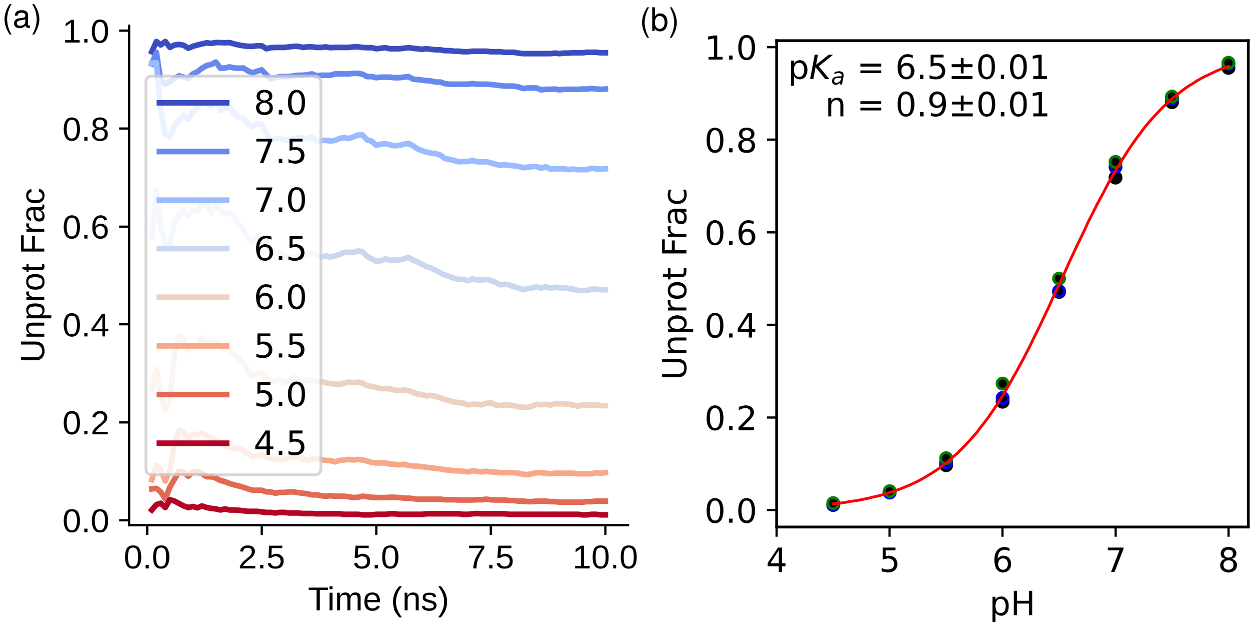


Figure S3: (a) Time series of the cumulative unprotonated fraction of His from the first set of three replica-exchange simulations. Legends indicate the pH conditions. (b) Unprotonated fractions of His at different pH. Data are from three independent sets of pH replica-exchange simulations of $CH_3COAAHAACONH_2$. The best fit (red) and the p$K_a$ shown were obtained using all data points. The p$K_a$ from the bootstrap method is 6.5±0.02.

.

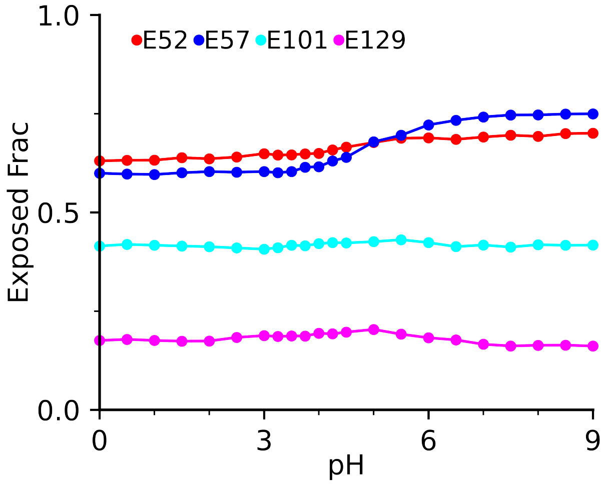


Figure S4: Fraction of SASA for E51, E57, E101 and E129 in SNase at different pH.

.

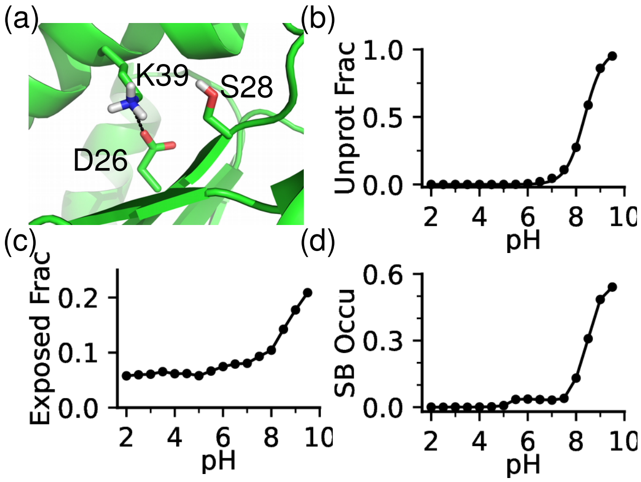


Figure S5: a) Zoomed-in view of the environment of Asp26 in thioredoxin taken from the simulation at pH 8. b) Unprotonation fraction of Asp26 at different pH. c) Fraction of solvent exposure of Asp26 side chain at different pH. c) Occupancy of the salt-bridge formation between Asp26 and Lys39 at different pH.

.

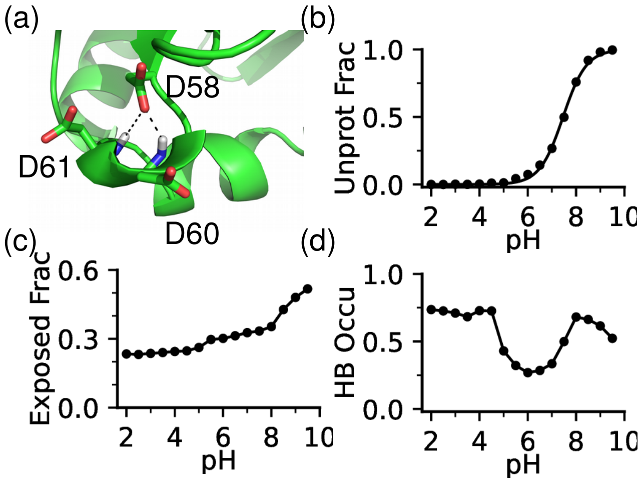


Figure S6: a) Zoomed-in view of the environment of Asp58 in thioredoxin taken from the simulation at pH 4. H-bond donors are the backbones of Asp61 and Asp60 . b) Unprotonation fraction of Asp58 at different pH. c) Fraction of solvent exposure of Asp58 side chain. c) Occupancy of the h-bond formation of Asp58 with Asp58 and Asp60.

.
